# Rab2 drives axonal transport of dense core vesicles and lysosomal organelles

**DOI:** 10.1101/2020.06.03.131672

**Authors:** Viktor K. Lund, Matthew D. Lycas, Anders Schack, Rita C. Andersen, Ulrik Gether, Ole Kjaerulff

## Abstract

Long range fast axonal transport of neuropeptide-containing dense core vesicles (DCVs), endolysosomal organelles and presynaptic components is critical for maintaining the functionality of neurons. How the transport of DCVs is orchestrated remains an important unresolved question. The small GTPase Rab2 has previously been shown to mediate DCV biogenesis and endosome-lysosome fusion. Here we use the *Drosophila* model system to demonstrate that Rab2 also plays a critical role in bidirectional axonal transport of DCVs, endosomes and lysosomal organelles, most likely by controlling molecular motors. We further show that the lysosomal motility factor Arl8 is required as well for axonal transport of DCVs, but unlike Rab2 is also critical for DCV exit from cell bodies into axons. Our results uncover the mechanisms responsible for axonal transport of DCVs and reveal surprising parallels between the regulation of DCVs and lysosomal motility.

## INTRODUCTION

Peptide neurotransmitters and hormones are critical for controlling behavior, metabolism, and growth. In neurons, neuropeptides released from both synaptic and extrasynaptic sites mediate direct neurotransmission, neuromodulation and endocrine functions. Neurons and secretory cells use secretory granules or dense core vesicles (hereafter referred to as DCVs) to store and transport neuropeptides and peptide hormones prior to regulated release. After synthesis in the endoplasmic reticulum (ER) and transit through the Golgi apparatus, neuropeptides and accessory proteins are packaged into immature DCVs that bud from the *trans*-Golgi network (TGN). This is followed by a maturation process involving exchange of material between immature DCVs and the endosomal system, whereby inappropriate proteins are removed while genuine DCV cargoes are retained (Gondre-Lewis et al., 2012; Topalidou et al., 2016).

Small GTPases of the Rab and Arf/Arl families regulate organelle identity, fusion and motility in eukaryotic cells (Donaldson and Jackson, 2011; Stenmark, 2009). They cycle between an inactive, GDP-bound, cytosolic state and an active, GTP-bound, membrane-associated state, in which they recruit specific effector proteins such as molecular tethers and motors to the membranes of their cognate organelles. In nematodes and mammals, the highly conserved Rab family GTPase Rab2 has an important role in DCV maturation. Together with its effector CCCP-1 and the endosome-associated recycling protein (EARP) tethering complex, Rab2 promotes retrieval of regulated secretory cargo from the endosomal system to maturing DCVs (Ailion et al., 2014; Laurent et al., 2018; Topalidou et al., 2016), and impairment of Rab2 or its effectors leads to the loss of DCV cargo to the endosomal system (Edwards et al., 2009; Sumakovic et al., 2009). Recently, we and others reported that Rab2 is critical for biogenesis of mature degradative endo- and autolysosomal compartments by controlling the fusion of late endosomes (LEs) with lysosomes and transport vesicles carrying lysosomal membrane proteins, and fusion of autophagosomes with endolysosomal organelles (Lorincz et al., 2017; Lund et al., 2018) (Ding et al., 2019).

Unlike small synaptic vesicles, neuronal DCVs cannot regenerate locally, and must be transported from the soma through axons (or dendrites) to their release sites. Generally, axonal transport relies on movement of organelles and protein complexes by molecular motors along axonal microtubules (MTs). Anterograde transport is mediated by the diverse family of plus-end-directed kinesin motors, while retrograde transport depends on cytoplasmic dynein. However, a high degree of interdependence exists between anterograde and retrograde transport (Barkus et al., 2008; Lim et al., 2017; Martin et al., 1999; Twelvetrees et al., 2016). Motor recruitment and regulation is mediated by adaptor proteins and small GTPases and is organelle-specific, producing different movement patterns for different organelles (Guedes-Dias and Holzbaur, 2019). In both mammals and invertebrates, axonal transport of DCVs is mediated by the Kinesin-3 KIF1A/unc-104 and Kinesin-1 (Barkus et al., 2008; Gumy et al., 2017; Lim et al., 2017; Lo et al., 2011), but how these motors are recruited and regulated is not well understood.

Axonal transport of endosomes and lysosomal organelles is also vital for neuronal development and function, partially through delivery of building blocks for axonal outgrowth and presynaptic biogenesis (Farias et al., 2017; Vukoja et al., 2018). Axonal transport of LEs and lysosomes critically depends on the BORC-Arl8 complex, which recruits Kinesin-1 or Kinesin-3 motors (Farias et al., 2017; Rosa-Ferreira and Munro, 2011; Wu et al., 2013). Perturbation of axonal transport can have devastating consequences for human health and has been implicated in a number of nervous system disorders, most notably neurodegenerative and neurodevelopmental conditions (De Vos et al., 2008; Guedes-Dias and Holzbaur, 2019).

Here we investigate the role of Rab2 in DCV biogenesis and transport in the *Drosophila* model system. We show that *Drosophila* Rab2 promotes DCV biogenesis and neuropeptide storage. Surprisingly, we also find that active Rab2 localizes to moving DCVs in axons and is required for fast and processive bidirectional axonal transport of DCVs, but not for DCV axonal entry from neuronal cell bodies or for synaptic capture and regulated exocytosis. Loss of Rab2 causes accumulation of static and slowly moving DCVs in the axonal compartment, and reduces the amount of DCVs at synaptic terminals. In line with this, Rab2 exhibits activity-dependent nanometer-range proximity to the Kinesin-3 unc-104 *in vivo*, suggesting that active Rab2 is part of a motor protein-containing molecular complex located on the surface of DCVs. We further find that the LE/lysosomal GTPase Arl8, but not Rab7, is also involved in axonal transport of DCVs. Unlike Rab2, however, Arl8 is both needed for axonal entry of DCVs from neuronal cell bodies and for their transport through the axon. Rab2 is also required for effective axonal transport of early endosomes (EEs), LEs, lysosomes, and lysosome-related active-zone transport vesicles. Our results demonstrate that the small GTPases Rab2 and Arl8 are essential regulators of axonal DCV transport, and reveal a novel role for Rab2 in the axonal transport of endosomes and lysosomes.

## RESULTS

### Rab2 is enriched in peptidergic neurons and required for DCV biogenesis

Detailed examination of the expression pattern of hemagglutinin-tagged Rab2 (HA-Rab2) controlled by endogenous *Rab2* regulatory elements (Lund et al., 2018) in third instar (L3) larval brains revealed that Rab2 is enriched in somata of long-axon high-capacity peptidergic neurons expressing the DIMM transcription factor (Hamanaka et al., 2010; Park et al., 2008) (Fig. 1A-C). This is consistent with *Rab2* being a transcriptional target of DIMM (Hadzic et al., 2015). Among the Rab2-enriched neurons were insulin-producing cells (IPCs) that synthesize and release insulin-like peptides (Ilp) 2, 3, and 5 to control growth and metabolism (Rulifson et al., 2002). A strong HA-Rab2 signal was detected in both IPC somata and axonal projections to the aorta and ring gland (Fig. 1D). Compared to wild type, the Ilp2 content in IPC somata in normally fed larvae was substantially reduced (to one third or less) in *Rab2*^*Δ1*^ null mutants (Fig. 1E, F) or when *Rab2* was depleted by RNA interference (RNAi) (Fig. S1A, B). The *Rab2*^*Δ1*^ phenotype in IPCs was rescued by expressing an *mCherry-Rab2* transgene (Fig. 1E, F). Strikingly, the starvation-induced accumulation of Ilp2 in IPC somata of wild type larvae (Fig. 1G) (Geminard et al., 2009) did not occur in *Rab2*^*Δ1*^ animals (Fig. 1G).

**Figure 1.**
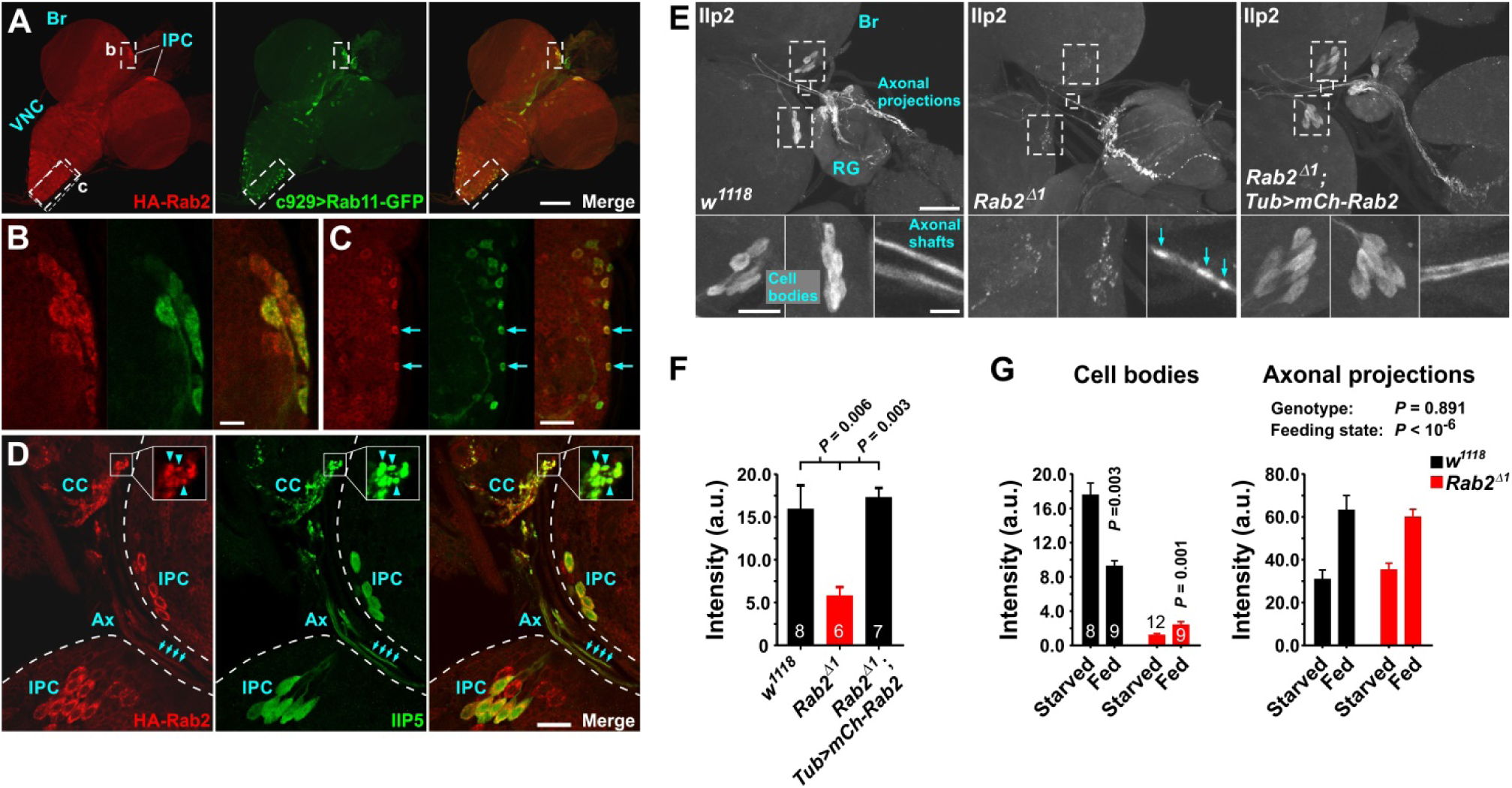
Rab2 is enriched in high-capacity peptidergic neurons in the *Drosophila* CNS and is required for Ilp2 accumulation in IPC cell bodies. **(A)** CNS from late L3 larva expressing *HA-Rab2* controlled by the endogenous *Rab2* promoter, and *c929*>GFP-Rab11 to mark DIMM-expressing peptidergic neurons. Br, brain hemispheres; VNC, ventral nerve cord; IPC, insulin-producing cells. Overlap between *c929*>GFP-Rab11 and HA-Rab2 immunostaining in IPC somata **(B)** and VNC peptidergic somata **(C)** (*arrows*) is shown. **(D)** Brain IPC somata (IPC) and axonal projections (Ax) to the dorsal aorta and corpora cardiaca (CC) in the ring gland of *HA-Rab2* larva immunostained for HA and Ilp5. Note presence of Rab2 in IPC somata, axonal shafts (*arrows*), and distal neurohemal release sites at the ring gland (*inset, arrowheads*). **(E)** Larval brain (Br) and ring gland (RG) in wild type, *Rab2*^*Δ1*^, and *Rab2*^*Δ1*^ rescued by expressing *mCherry-Rab2* under the *Tubulin* promoter, immunostained for Ilp2. **(F)** Ilp2 immunosignal intensity in IPC somata in **(E)**. ANOVA followed by Tukey’s test. **(G)** Ilp2 intensity in larval IPC cell bodies and axonal projections. Two-factor ANOVA, at *left* followed by Tukey’s test. Scale bars: **(A)**, 100 µm; **(B)**, 10 µm; **(C)**, 10 µm; **(D)**, 20 µm; **(E)** *top*, 50 µm; *bottom left* and *middle insets*, 20 µm; *bottom right inset*, 5 µm. In **(F)** and **(G)**, data are mean ± s.e.m., and the number of preparations analyzed for each genotype is indicated *inside bars*. Source data underlying this and the subsequent Figures are provided as a Source Data file, and complete genotypes are listed in Table S1.

The reduction in Ilp2 content in *Rab2*^*Δ1*^ mutants was not due to post-transcriptional effects, as similar decreases in IPC cell body content were observed in *Rab2* nulls for GFP-tagged preproAtrial Natriuretic Factor (ANF-GFP), a DCV luminal cargo reporter (Rao et al., 2001), and for a GFP-tagged variant of the DCV transmembrane protein IA-2 (IA2-GFP) when expressed in peptidergic cells driven by *386Y*-Gal4 (Fig. S2A, B, D, E).

Microscopy and western blotting revealed that *Rab2*^*Δ1*^ larvae exhibited a substantial reduction in whole-central nervous system (CNS) content of ANF-GFP expressed generally in peptidergic cells (*386Y*>ANF-GFP) (Fig. S3A-C). We also observed a marked increase in the amount of a lower molecular weight ANF-GFP cleavage product in *Rab2*^*Δ1*^ CNS samples (Fig. S3C), indicating that loss of Rab2 may alter neuropeptide processing. No changes were observed upon *Rab2* overexpression (Fig. S3C), suggesting that Rab2 is not rate limiting for neuropeptide storage or processing in wild type. Moreover, direct stochastic optical reconstruction microscopy (*d*STORM) demonstrated a small increase in DCV size in *Rab2*^*Δ1*^ specimens (Fig. S3E, F), consistent with DCV size alterations commonly associated with defects in DCV biogenesis (Asensio et al., 2013; Grabner et al., 2006; Sumakovic et al., 2009). In large peptidergic somata HA-Rab2 tightly enveloped TGN clusters (Fig. S3G) where DCV biogenesis occurs (Gondre-Lewis et al., 2012). Furthermore, RNAi depletion of Rab2 strongly reduced IPC levels of the DCV biogenesis- and growth-promoting PICK1 protein (Fig. S1A, C), which is bound and stabilized by the Rab2 effector ICA69 (Cao et al., 2013; Herlo et al., 2018; Holst et al., 2013). However, genetic depletion of PICK1 or the Rab2 effectors *CG3703* (homolog of RUND-1) and Golgin104 (homolog of CCCP-1), implicated in DCV biogenesis in nematodes (Ailion et al., 2014), or the Rab2 effectors Golgin-45, BicD, and Vps39 (Gillingham et al., 2014) involved more generally in intracellular trafficking failed to fully recapitulate the *Rab2*^*Δ1*^ neuropeptide storage phenotype (Fig. S3C, D). Strong hypomorphic mutants for *Vps41/light* (*lt*) and *Vps18*/*deep orange* (*dor*), additional components of the HOPS complex mediating Rab2-dependent in LE-lysosome fusion, also did not exhibit reductions in IPC cell body Ilp2 content (Fig. S3D).

Consistent with an important role of Rab2 in neuropeptidergic function, RNAi depletion of Rab2 specifically in peptidergic neurons resulted in either death prior to eclosion, or wing expansion defects (Fig. S4A-C) associated with dysfunction of the protein hormone Bursicon (Dewey et al., 2004), secreted from DIMM-positive (Park et al., 2008), Rab2-enriched neurons (Fig. S4D).

These findings indicate that *Drosophila* Rab2 is upregulated in peptidergic neurons and is involved in DCV biogenesis.

### Rab2 is required for DCV axonal transport

A striking feature of the *Rab2*^*Δ1*^ phenotype in larval brain IPCs was that, despite the dramatic reduction in Ilp2 somata, the Ilp2 signal was not reduced in the projections to the dorsal aorta and ring gland (Fig. 1E, G). This was observed in both starved and fed larvae (Fig. 1G), and for ANF-GFP expressed in IPCs (Fig. S2A-C). Furthermore, the IA2-GFP signal was substantially increased in the aortal projection of *Rab2*^*Δ1*^ larvae (Fig. S2D, F), despite a strong IA2-GFP reduction in IPC somata (Fig. S2D, E). Also, we observed focal Ilp2 accumulations in axonal shafts of *Rab2*^*Δ1*^ IPCs, but not in wild type or rescued specimens (Fig. 1E, lower right insets). Similar results were obtained in Bursicon-containing neurons in the larval ventral nerve cord (VNC) that project commissural axons across the VNC midline, where a collateral branches off in each segment to form a local arbor (Fig. 2A, asterisks). In *Rab2*^*Δ1*^ larvae, the Bursicon content was strongly elevated in both the midline arbors and in focal accumulations in the commissural axonal shafts (Fig. 2A, B). These observations indicate that neuropeptides suffer an axonal transport defect in *Rab2*^*Δ1*^ animals.

**Figure 2.**
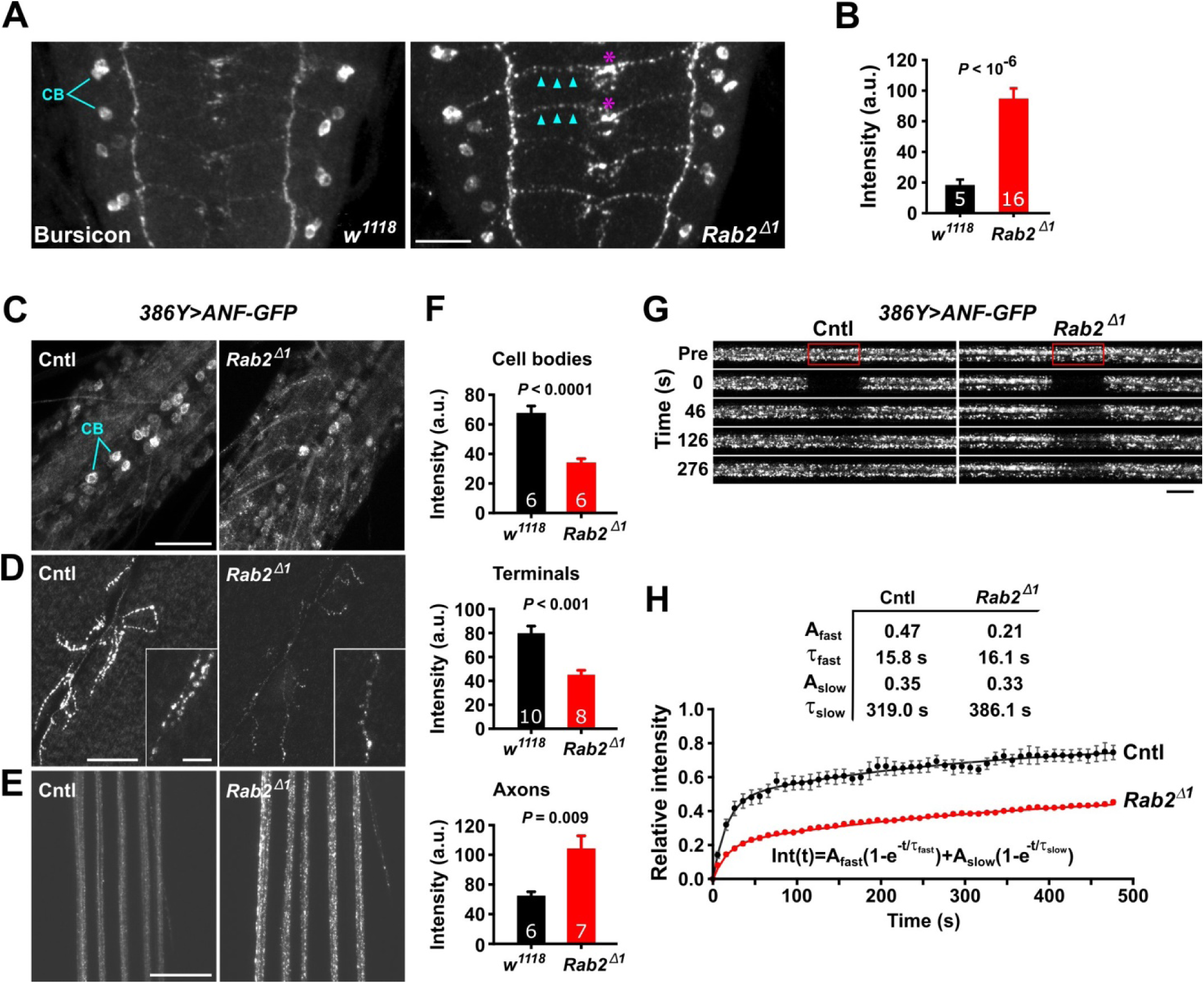
Static DCVs accumulate in axons in *Rab2*^*Δ1*^ larvae. **(A and B)** Increased bursicon levels in *Rab2*^*Δ1*^ neurites. **(A)** Anti-bursicon immunostaining in larval VNC. Dorsal view of 4 abdominal segments. Anterior at the *top*. Cell bodies (CB) are indicated. **(B)** Bursicon immunosignal in midline arbors in **(A)**. Student’s *t*-test. A.u., arbitrary units. **(C-E)** Sum projections of living fillet-dissected control and *Rab2*^*Δ1*^ larvae expressing ANF-GFP in peptidergic neurons driven by *386Y*-Gal4. Cell bodies (CB) in the dorsal VNC are indicated in **(C). (D)** Synaptic terminals of the muscle 6/7 NMJ. **(E)** Peripheral nerves proximal to the VNC. **(F)** Quantification of **(C-E)**. Student’s *t*-tests. **(G)** FRAP experiments revealing the motility of ANF-positive DCVs in axons of A8/9 nerves. DCVs imaged before (Pre) and at the indicated time points after photobleaching the central bleach box (*red*). **(H)** Data from **(G)** normalized to pre-bleach signal intensity and adjusted for bleaching through imaging (see Methods). Error bars are s.e.m.. Double exponential fits to data from eight separate experiments in five control larvae, and seven in five *Rab2*^*Δ1*^ larvae. In **(B)** and **(F)**, data are mean ± s.e.m., and *numbers inside bars* indicate the number of preparations analyzed for each genotype. Scale bars: **(A)**, 20 µm; **(C)**, 50 µm; **(D)**, 40 µm, (*inset*, 10 µm); **(E)**, 40 µm, **(G)**, 10 µm.

Live confocal imaging of peripheral nerves of filleted *386Y*>*ANF-GFP* L3 larvae revealed a substantial accumulation of ANF-GFP DCV cargo in axons of *Rab2*^*Δ1*^ peptidergic neurons (Fig. 2E). In contrast, ANF-GFP cargo in peptidergic somata in the VNC (Fig. 2C) and muscle 6/7 NMJ terminals (Fig. 2D) in living animals, and in 1b NMJ terminals in muscle 3 in fixed larvae (Fig. S5A, B) was reduced in *Rab2*^*Δ1*^ mutants. Fluorescence recovery after photobleaching (FRAP) in A8/9 peripheral nerves showed that ∼53% of the DCV material in wild type nerves exchanged slowly (t_1/2_ > 200 s) or remained static, while in *Rab2*^*Δ1*^ nerves this parameter increased to 79 % (Fig. 2G, H). These findings suggest that without Rab2, DCVs stall in the axonal compartment.

To investigate DCV transport parameters in more detail, we used GFP-tagged Ilp2 (ILP-GFP) (Wong et al., 2012) as a DCV cargo marker, controlling its expression with the motor neuron driver *OK6*-Gal4. The Ilp2-GFP signal is substantially brighter than ANF-GFP, and examining motor neurons allowed us to differentiate between anterograde and retrograde transport. In *Rab2*^*Δ1*^ larvae, focal accumulations of Ilp2-GFP appeared in axons near the roots of A1-7 nerves and along much of the A8/9 nerves. We therefore imaged the A7 nerve 0.5-1.0 mm from the VNC to avoid potential steric (traffic jam type) effects on transport. We photobleached a 60 µm long nerve segment on each side of a 10 µm central region, so that DCVs initially located in the unbleached center could be monitored with limited interference from unbleached DCVs entering from the sides (Fig. 3A, Video 1). While in *Rab2*^*Δ1*^ mutants no DCV cargo accumulated in A7 motor axons (Fig. S6B, C), the proportion of static DCVs approximately tripled (Fig. 3A, B, Video 1), and DCV speed during both anterograde and retrograde runs (i.e., bouts of uninterrupted movement at constant velocity) was reduced (Fig. 3D). Moreover, pause durations and the relative time spent pausing (duty cycle) were substantially increased for dynamic DCVs in *Rab2*^*Δ1*^ (Fig. 3E, F). As reported earlier (Barkus et al., 2008), most moving vesicles exhibited a preferred direction, either anterograde or retrograde, punctuated by pauses or shorter runs in the opposite direction. To estimate the processivity (length of uninterrupted movement between pauses) of axonal DCV transport, we divided the total run length in a particular direction by the number of pauses preceded by a run in that direction. The mean distance of uninterrupted movement in wild type - 73.7 ± 5.7 µm (anterograde), 44.4 ± 2.1 µm (retrograde) - was significantly reduced to 42.8 ± 3.1 and 24.0 ± 3.1 µm, respectively, in *Rab2*^*Δ1*^ animals (Fig. S6A). Furthermore, in *Rab2*^*Δ1*^ nerves markedly fewer DCVs traveled in both anterograde and retrograde directions, although retrograde transport was more severely affected (Fig. 3A, B, Video 1). Similar reductions in DCV flux were detected in single distal axons supplying muscle 3 Ib terminals (Fig. 4A, Fig. S5C, E). The *Rab2*^*Δ1*^ transport phenotype in peripheral nerves was rescued by expressing *Rab2* specifically in motor neurons (Fig. 3B-F, Fig. S6A and Video 1) confirming that it is cell-autonomous. Notably, *OK6*-Gal4-driven expression of *Rab2* partially restored adult viability in *Rab2*^*Δ1*^ animals, although the emerged adult flies suffered from wing expansion defects and died shortly after eclosion.

**Figure 3.**
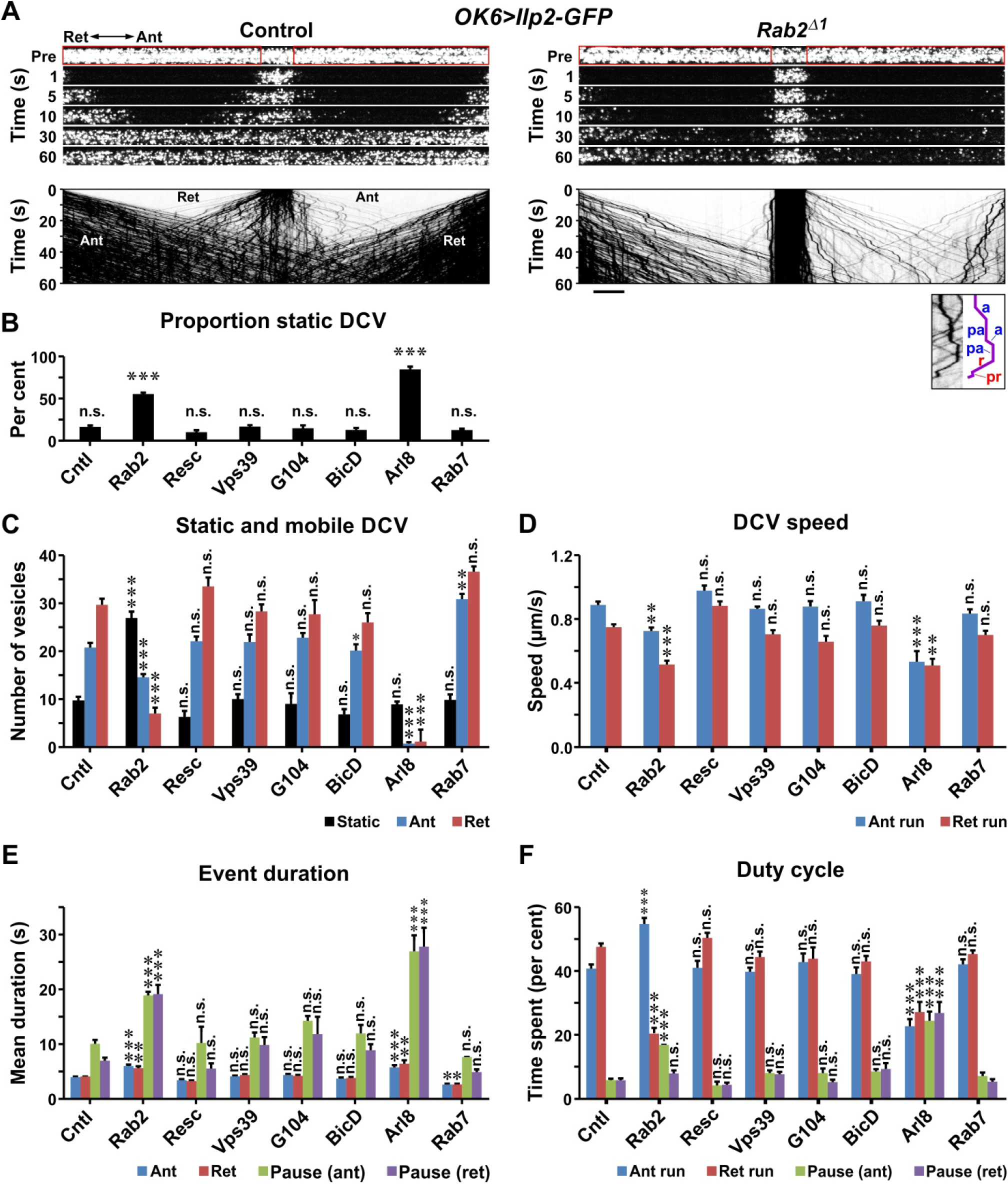
Loss of Rab2 disrupts axonal DCV transport. **(A)** (*top*) Time-lapse imaging showing transport of Ilp2-positive DCVs in motor axons of A7 nerves. DCVs imaged before (Pre) and at the indicated time points after photobleaching the lateral areas (*red boxes*). Ant, anterograde; Ret, retrograde. **(A)** (*bottom*) Kymographs (inverted images) derived from the time-lapse series in **(A)** (*top*). The image detail/diagram at the *lower right* depicts transport event categories: anterograde run (a), retrograde run (r), pause preceded by anterograde run (pa), pause preceded by retrograde run (pr). **(B)** Proportions of static vesicles, derived from **(C). (C)** Frequency distribution of static or mobile DCVs. Note the severely reduced number of vesicles in Arl8. **(D-F)** Kinetic parameters of mobile DCVs. In **(B-F)** data are mean ± s.e.m, and genotypes (number of preparations) are: Control (27), *Rab2*^*Δ1*^ (21), *Rab2*^*Δ1*^; *UAS-Rab2/+* (7), *Arl8*^*e00336*^*/Df* (14), *Vps39*^*Δ1*^*/Df* (5), *G104*^*ins1*^*/Df* (5), *BicD*^*r5*^*/Df* (5), *Rab7*^*Δ1*^ (6). All genotypes also included *OK6*>*Ilp2-GFP*. ANOVAs followed by Dunnett’s test. \**P*<0.05; \*\**P*<0.0001;\*\*\**P*<0.0000001. N.s., not significant. Scale bar, 10 µm.

**Figure 4.**
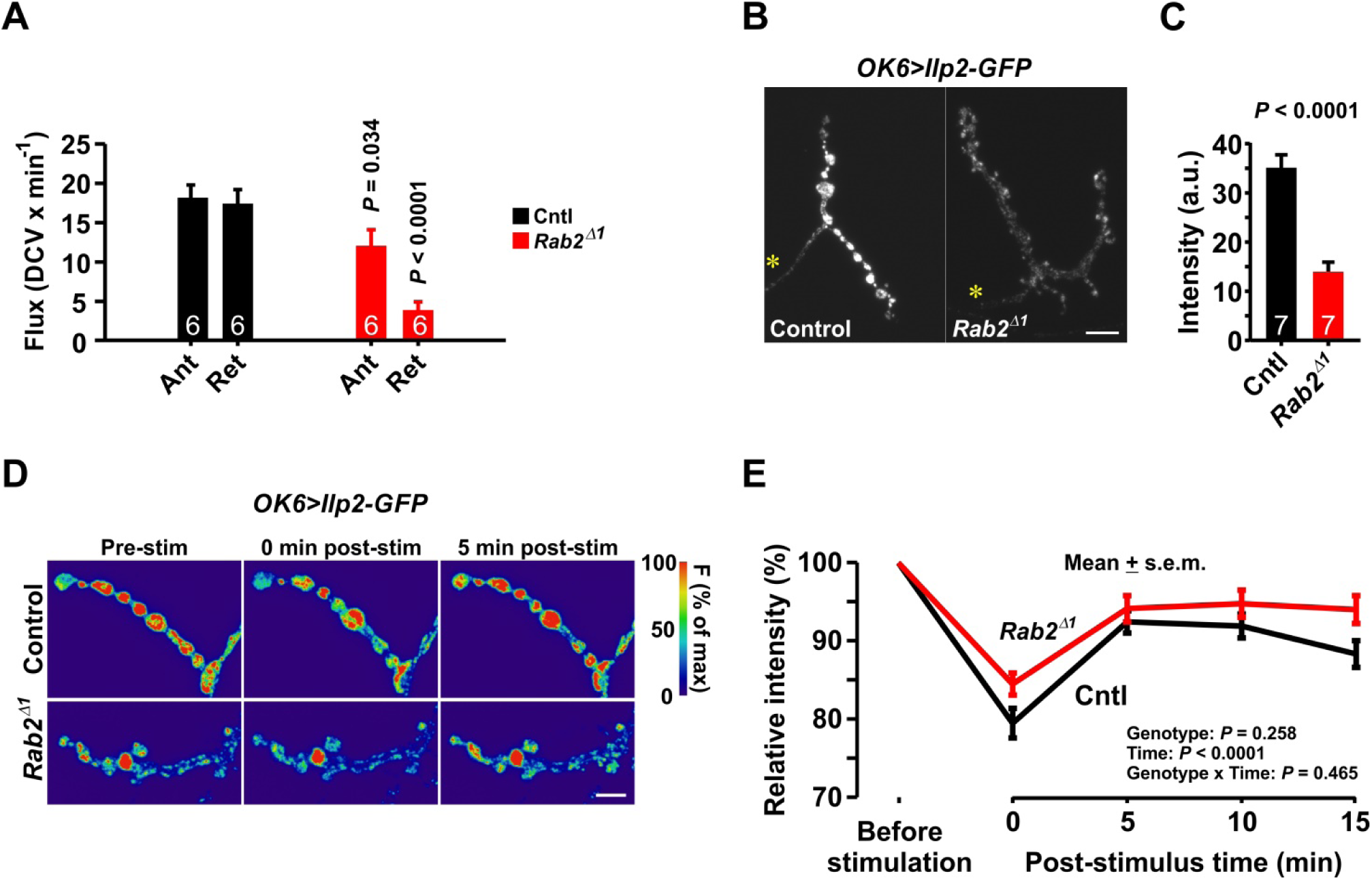
Rab2 loss reduces DCV flux in the distal axon, and the presynaptic resting level of Ilp2-GFP at NMJs, but leaves stimulated Ilp2-GFP release and replenishment largely unaffected. **(A)** Flux of Ilp2-GFP-positive DCVs transported in the distal axon and crossing near the *yellow asterisks* in **(B)**. Ant, anterograde; Ret, retrograde. Student’s *t*-tests. **(B)** Sum projections of the muscle 3 NMJ in larvae expressing Ilp2-GFP in motor neurons (live imaging). **(C)** Quantification of **(B)**. Student’s *t*-test. **(D)** False-color coded GFP fluorescence intensity before, immediately after, and five minutes after high K^+^-induced Ilp2-GFP release. For each genotype, pixel values were normalized to the maximum intensity across images (100%, red). **(E)** GFP fluorescence relative to the pre-stimulus level. Two-factor ANOVA. Data from 14 control and 13 *Rab2*^*Δ1*^ larvae. In **(A)** and **(C)**, data are mean ± s.e.m., and the number of larvae analyzed is indicated *inside bars*. Scale bars: **(B)**, 10 µm; **(D)**, 5 µm.

Comparable results were obtained when imaging transport of ANF-GFP-positive DCVs in A8/9 nerves of *386Y*>*ANF-GFP* larvae, although we could not differentiate between anterograde and retrograde transport due to the mixed orientation of the involved axons (Fig. S5F-J).

When expressing Ilp2-GFP in pharate adult stage Bursicon-secreting neurons, and depleting Rab2 using RNAi, the axonal transport defect observed in *Rab2*^*Δ1*^ larval motor neurons was essentially reproduced (Fig. S7A-H). A subset of these neurons, located in the VNC, project axons densely adorned with neurohormone-filled *en passant* boutons into the body cavity (Fig. S7A). We quantified DCV transport in single axonal shafts at a point near the VNC (Fig. S7A, yellow asterisk). *Rab2* knockdown strongly reduced bidirectional flux, preferentially affecting the retrograde component (Fig. S7E, F). A reduction of the anterograde speed of transiting DCVs (Fig. S7G) and a large increase in the proportion of static DCVs in the axon (Fig. S7H) was also observed. Overall, the Ilp2-GFP signal in the bouton-containing distal axons was reduced in Rab2-depleted neurons (Fig. S7A, B, D), whereas aberrant focal accumulations of DCV material were readily apparent in the more proximal axonal shafts in the dorsal nerve (Fig. S7B). Interestingly, Rab2 overexpression resulted in a depletion of DCV signal in the bouton-containing distal axons and a simultaneous increase in the cell bodies, without reducing anterograde axonal flux (Fig. S7A, C-F), indicating that Rab2 may promote retrograde DCV transport.

Staining for endogenous Rab7, and live imaging of mCherry-Rab7, showed either no or only a modest overlap between LEs and ANF-GFP-marked DCVs in mid-nerve axonal shafts of *Rab2* null larvae (Fig. S6D, E and Video 2). Therefore, the apparent alterations in DCV axonal transport are not caused by rerouting of DCV cargo to LEs, which are expected to have different transport characteristics than *bona fide* DCVs.

Importantly, *Vps39*^*Δ1*^ hemizygous null mutant larvae, in which LE-lysosome fusion is blocked as in *Rab2* nulls (Lorincz et al., 2017; Lund et al., 2018), did not display the defects in transport of Ilp2-GFP-positive DCVs in motor axons that characterized *Rab2*^*Δ1*^ animals (Fig. 3B-E). Likewise, mutating the genes encoding the Rab2 effectors Golgin104 and BicD did not affect DCV transport (Fig. 3B-E). Similarly, loss of these three Rab2 effectors did not recapitulate the *Rab2*^*Δ1*^-associated DCV transport phenotype in peripheral nerve axons of *386Y*>*ANF-GFP* larvae (Fig. S5G-J). Therefore, the perturbed axonal transport in *Rab2* nulls is not due to a block in the endo-lysosomal and autophagic pathways and also cannot be explained by the dysfunction of the DCV maturation process controlled by Rab2 through the Golgin104/CCCP-1 pathway. Moreover, the Rab2-binding dynein adaptor BicD does not play a major role in axonal transport of DCVs.

These data show that Rab2 is required for normal axonal transport of DCVs, but not for DCV entry from the cell body into the axon.

### Rab2 is not important for stimulated release and synaptic capture of DCVs

Since Rab2 is involved in sorting of DCV components during DCV biogenesis (Ailion et al., 2014; Edwards et al., 2009; Sumakovic et al., 2009; Topalidou et al., 2016), we tested if Rab2 is required for stimulated exocytosis of DCV cargo. We stimulated *OK6*>*Ilp2-GFP* larval fillets by K^+^-induced depolarization and monitored both DCV cargo release and subsequent replenishment in muscle 3 type Ib NMJs as changes in synaptic Ilp2-GFP fluorescence. As for ANF-GFP (Fig. S5A, B), the Ilp2-GFP content of muscle 3 type 1b terminals of *Rab2*^*Δ1*^ mutants was substantially reduced (Fig. 4B, C), likely due to decreased anterograde DCV flux in the distal axon (Fig. 4A). Also, terminals often appeared morphologically aberrant (Fig. 4B and Fig S5A) (Mallik et al., 2017). However, both the proportional DCV cargo release upon stimulation and the subsequent partial recovery were similar in mutant and wild type terminals, although the relative release amplitude tended to be somewhat lower in *Rab2*^*Δ1*^ (Fig. 4D, E). Hence, Rab2 is neither critically required for stimulated exocytosis of DCVs, nor for the fast post-stimulus recovery of the synaptic DCV pool that relies on activity-induced capture of anterograde circulating DCVs (Cavolo et al., 2016; Shakiryanova et al., 2006).

Without stimulation, DCVs circulate between the axon initial segment and terminal boutons, with a low probability of being captured when passing through individual *en passant* boutons (Wong et al., 2012). In *Rab2*^*Δ1*^ larvae, Ilp2-GFP-positive DCVs in both *en passant* and terminal boutons of muscle 3 type Ib terminals did not display increased synaptic capture (Fig. S8A-C) or decreased retrograde movement from the terminal bouton (Fig. S8D-F), suggesting that the increase in static DCVs in axonal shafts and the deficit in retrograde DCV flux in *Rab2* nulls is not due to enhanced baseline DCV capture. Furthermore, the single-vesicle Ilp2-GFP intensity was preserved in *Rab2* nulls, suggesting that the decreased DCV cargo content in *Rab2*^*Δ1*^ mutant terminals is due to lower vesicle (quantal) content in boutons rather than reduced cargo loading in individual vesicles (Fig. S8G).

These findings show that Rab2 is neither critically required for regulated exocytosis of DCVs, nor does it play a major role in synaptic capture.

### Rab2 controls axonal transport of endosomes and lysosomal organelles, but not mitochondria

To test if Rab2 exclusively regulates axonal transport of DCVs, we examined the distribution and motile characteristics of mitochondria and organelles belonging to the endolysosomal family. We detected no difference in axonal transport of mito-GFP-labeled mitochondria between *Rab2*^*Δ1*^ and wild type controls (Fig. S9A, B). Also, the axonal tubulin cytoskeleton was unperturbed when visualized by immunolabeling the axonal microtubule-binding protein Futsch (Hummel et al., 2000), both in terms of staining intensity and microtubule continuity in the distal axon and presynaptic arbor of muscle 3 1b terminals (Fig. S9C).

Rab2 is closely associated with the function of the endolysosomal system and is present on the membranes of endosomes and lysosomes, albeit transiently or in very low abundance in most cell types (Gillingham et al., 2014; Lorincz et al., 2017; Lund et al., 2018). Accordingly, both EEs, marked by GFP-Rab5 and LEs, marked by GFP-Rab7, showed markedly reduced motility in peripheral motor axons in the absence of Rab2. Wild type axons exhibited both a brightly labeled static and a considerable, but more weakly labeled, dynamic population of EEs undergoing bidirectional transport (Fig. 5A, Video 3). In *Rab2*^*Δ1*^ larvae, both the anterograde and retrograde dynamic EEs were almost entirely absent, while the proportion of static GFP-Rab5-positive organelles was strongly increased (Fig. 5A, Video 3). A large population of motile LEs was also present in wild type axons, in addition to a minor static component. In contrast to EEs, however, LEs in controls exhibited a significant directional bias towards retrograde transport (Fig. 5B, Video 4), as described in other systems (Cheng et al., 2015; Ferguson, 2018). Rab2 loss increased the proportion of static LEs, decreased the number and average velocity of motile LEs, and strongly reduced the retrograde transport bias (Fig. 5B, Video 4). However, the overall effect was less dramatic than for EEs.

**Figure 5.**
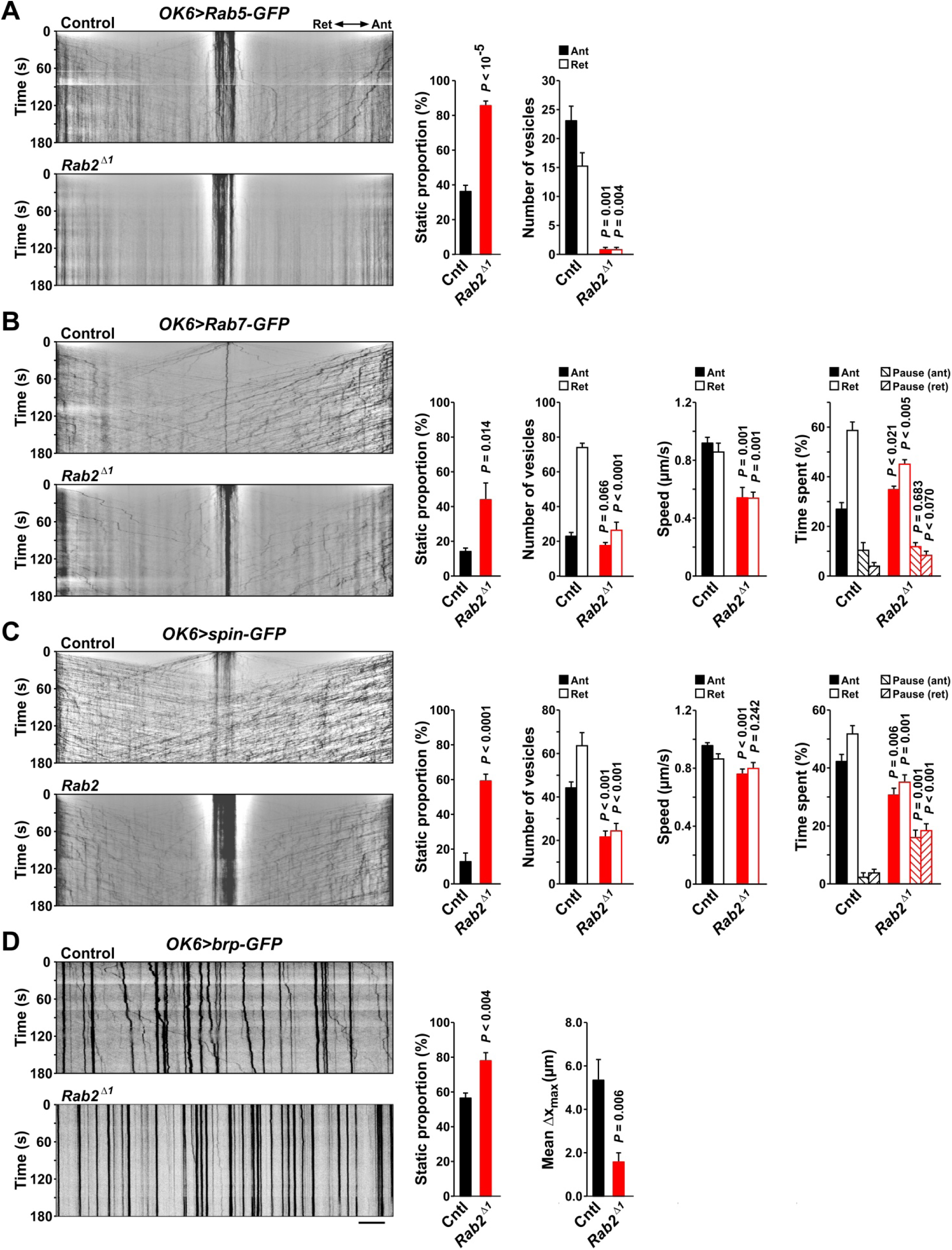
Rab2 loss strongly reduces the axonal motility of organelles associated with the endo-lysosomal system. Larval motor neurons expressing the EE marker Rab5-GFP **(A)**, the LE marker Rab7-GFP **(B)**, the lysosomal marker Spinster-GFP **(C)**, or the active zone-protein Bruchpilot carried in lysosome-related presynaptic transport vesicles **(D)**, all driven by *OK6-*Gal4. Kymographs (inverted images) depicting the transport of GFP-labeled organelles are shown at the *left* in **(A-D)**. In **(A-C)** the imaged field was initially photobleached bilaterally, as in (Fig. 3). To the *right* of the kymographs is shown the proportion of static organelles **(A-D)**, the organellar flux in a period of 180 s **(A-C)**, and the run speed and duty cycle **(B and C). (D)** (*right*) The maximal lateral excursion of brp-GFP-positive organelles. In each panel **(A-D)**, data are mean ± s.e.m., with 5 to 7 larvae included per genotype. The exact sample size is available in the Source Data file. Student’s *t*-tests. The kymographs in **(A-C)** were high-pass filtered (see Methods). Scale bar, 10 µm.

Next, we examined the axonal motility of the lysosomal transmembrane marker Spinster-GFP (spin-GFP). Wild type axons contained a very high density of spin-GFP-positive lysosomes and lysosome-related vesicles undergoing bidirectional transport (Fig. 5C, Video 5). Similar to the results for endosomes and DCVs, loss of Rab2 led to a large increase in the abundance of static lysosomal organelles, accompanied by a striking build-up of spin-GFP signal in the axons (Video 5), as well as marked reductions in both counts and dynamic parameters of motile lysosomal organelles (Fig. 5C, Video 5).

Presynaptic components including active zone proteins are delivered to the synapse in non-degradative lysosome-related vesicles positive for transmembrane lysosomal markers such as Spinster (Vukoja et al., 2018). We tested if axonal transport of a GFP-tagged variant of the active zone scaffold Bruchpilot (brp-GFP) is perturbed in *Rab2*^*Δ1*^ larvae. In wild type L3 motor axons, relatively few brp-positive organelles were present. Those that were motile (43.5 %) exhibited a pronounced anterograde bias, consistent with their role in presynaptic biogenesis (Fig. 5D). Loss of Rab2 substantially increased the proportion of static brp-organelles and markedly reduced the mobility of the remaining dynamic organelles, but did not affect the anterograde transport bias (Fig. 5D).

These results show that, while Rab2 is not universally required for axonal transport, it regulates axonal transport of both DCVs and organelles belonging to the endolysosomal family, including specialized derivatives responsible for delivery of presynaptic components.

### Rab2 is present on moving DCVs

If Rab2 is directly involved in axonal transport of DCVs and endolysosomal organelles, either as a motor adaptor or as a regulator of other factors coupling to motors or microtubules, one would expect it to be present in the axono-synaptic domain and more specifically on the membranes of the transported organelles. HA-Rab2 was highly abundant in IPC axons and neurohemal release zones that harbor many Ilp2-containing DCVs (Fig. 1D) and was likewise present in axonal shafts located in larval peripheral nerves (data not shown). However, with standard confocal live imaging GFP-Rab2 appeared to be largely diffusely distributed in peripheral nerves, with occasional strongly labeled immobile organelles. We reasoned that if Rab2 is present on the membranes of moving DCVs it might be in very low abundance or very transiently, similar to its fleeting interaction with endosomal membranes (Lorincz et al., 2017; Lund et al., 2018). Moreover, its presence on the DCVs would be obscured by cytosolic Rab2 in the axoplasm and in the surrounding glial sheath. To visualize such low-level recruitment, we bleached a segment of peripheral nerve expressing fluorescently labeled Rab2 to eliminate signal from unbound cytosolic protein. This was followed by confocal imaging with a high pixel dwell time and a high degree of averaging to detect small amounts of Rab2 on motor-driven organelles entering the bleached region before diffusion of cytosolic material could “catch up”. Due to the slow scanning speed, moving vesicular organelles appeared as streaks with slopes inversely proportional to the velocity of movement, similar to kymographic representations (Fig. 6A). We observed a high density of GFP-Rab2-positive vesicles moving in axons with speeds of up to ∼3 µm/s (Fig. 6A, top). Similar motile vesicles were also brightly labeled with neuronally expressed gain-of-function GTP-locked pHluorin-Rab2^Q65L^ (Fig. 6A, bottom) or wild type pHluorin-Rab2 (Fig. 6B). In contrast, very few moving vesicles were labeled with inactive GDP-locked pHluorin-Rab2^S20N^ (Fig. 6A, middle), demonstrating that recruitment of Rab2 to organelles undergoing axonal transport depends on its activation state. Importantly, a sizable fraction of the moving GFP-Rab2-, pHluorin-Rab2-, and pHluorin-Rab2^Q65L^-positive organelles were ANF-mCherry-labeled DCVs (Fig. 6B). These results demonstrate that active Rab2 is present on DCVs undergoing axonal transport.

**Figure 6.**
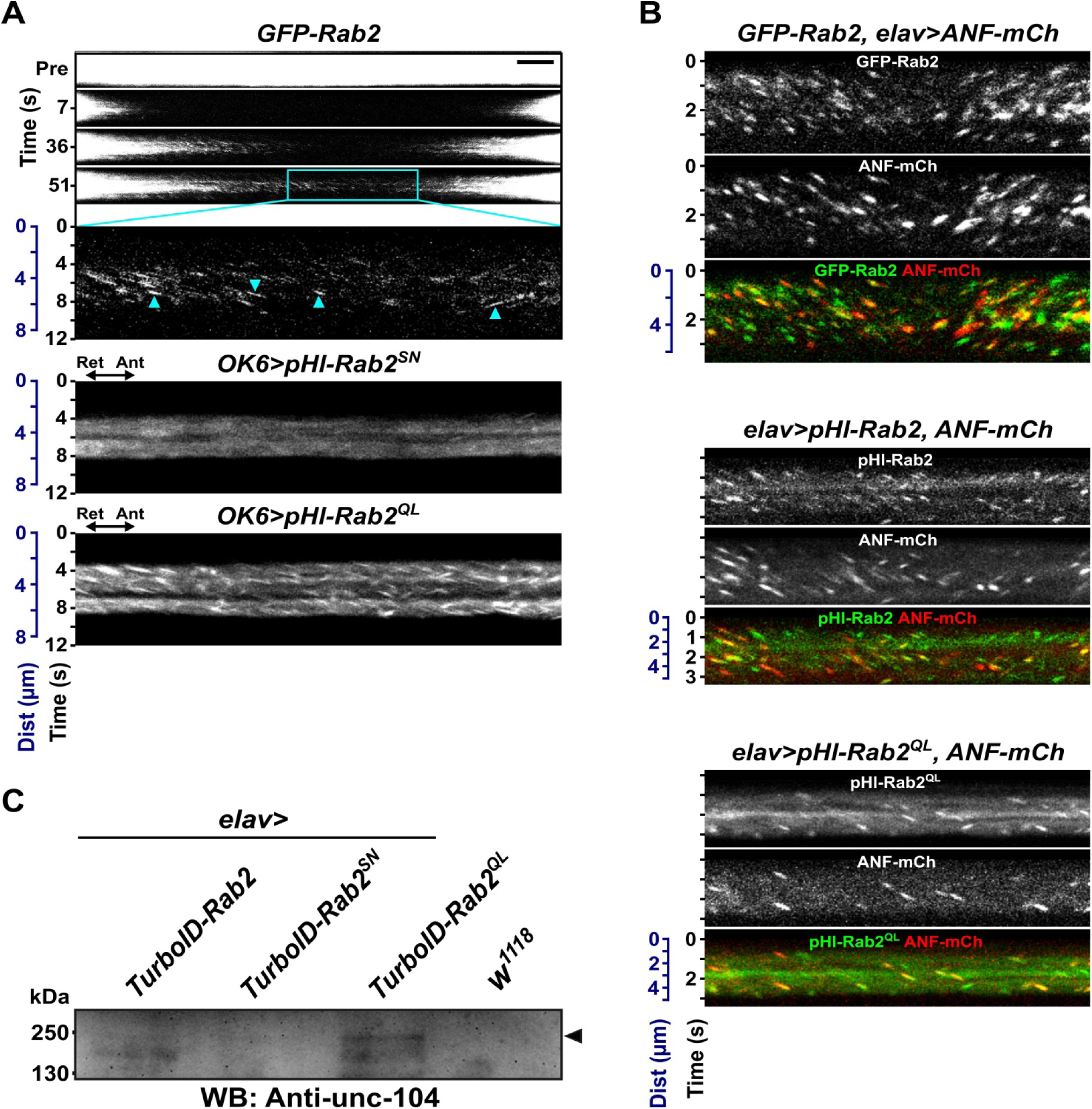
Active Rab2 is present on axonally transported DCVs and associates with Kinesin-3/unc-104 in the nervous system *in situ*. **(A)** (*top*) Axonal transport of GFP-Rab2-positive organelles. Fluorescence from cytosolic GFP-Rab2 and immobile structures was photobleached, followed by slow confocal scanning producing a kymograph-like image with the vertical dimension representing both distance and time (note double axis). Actively transported organelles invading from the flanking regions appear as streaks (*arrowheads*) with slopes inversely proportional to transport velocity. Also shown is inactive GDP-locked pHlourin-Rab2^S20N^ **(A)** (*middle*) and gain-of-function GTP-locked pHlourin-Rab2^Q65L^ **(A)** (*bottom*) in motor axons visualized as in **(A) (***top*). **(B)** Co-localization of GFP-Rab2 (*top*), pHlourin-Rab2 (*middle*), and pHlourin-Rab2^Q65L^ (*bottom*) with the DCV cargo marker ANF-mCherry in organelles undergoing axonal transport. Same imaging procedure as in **(A). (C)** Biotinylated proteins were purified from newly eclosed adults pan-neuronally expressing TurboID fused to Rab2, Rab2^S20N^, or Rab2^Q65L^, and eluates were immunoblotted against unc-104 (*lanes 1-3*). The *arrowhead* indicates the specific unc-104 band. Eluate from *w*^*1118*^ control flies not expressing TurboID is also included (*lane 4*). Dist, distance. WB, western blot. Images represent the following number of experiments (larvae): **(A)** *top*, 11(5); *middle*, 6(4); *bottom*, 6(3); **(B)** *top*, 14(4); *middle*, 5(1); *bottom*, 20(5). The western blot in **(C)** was replicated twice.

### Rab2 exhibits activity-dependent nanometer-range proximity to unc-104

To detect if active Rab2 forms a molecular complex with motor proteins responsible for axonal transport of DCVs *in vivo*, we employed proximity-dependent biotinylation. This approach relies on a promiscuous biotin ligase conjugated to a bait protein to biotinylate closely neighboring prey proteins, which can then be purified and detected. With the original BirA ligase this technique has a documented labeling radius of ∼10 nm (Kim et al., 2014) and is advantageous over classical co-affinity purification by allowing for detection of transient or weak molecular interactions. We found that a neuronally expressed fusion of the enhanced BirA variant TurboID (Branon et al., 2018) to gain-of-function Rab2^Q65L^ (TurboID-Rab2^Q65L^) biotinylates the main anterograde *Drosophila* DCV motor (Barkus et al., 2008; Lim et al., 2017), the Kinesin-3 unc-104 (Fig. 6C). In contrast, very little biotinylation of unc-104 was observed using wild type TurboID-Rab2, and essentially no biotinylation was observed using loss-of-function TurboID-Rab2^S20N^ or in control animals not carrying any transgenes (Fig. 6C). These results suggest that active Rab2 is part of a molecular complex that includes the anterograde DCV motor unc-104.

### Arl8, but not Rab7, is required for axonal transport of DCVs

The finding that the small GTPase Rab2 regulates biogenesis and axonal transport of both endolysosomal vesicles and DCVs may imply a connection between these organelles. This prompted us to ask if other small GTPases involved in the function of late-stage compartments of the endolysosomal pathway also play a role in axonal transport of DCVs. Apart from Rab2, fusion and microtubule-based motility of LEs and lysosomes is controlled by the small GTPases Rab7 and Arl8 (Boda et al., 2019; Farias et al., 2017; Johansson et al., 2007; Jordens et al., 2001; Khatter et al., 2015; Marwaha et al., 2017; Pankiv et al., 2010; Raiborg et al., 2015; Rosa-Ferreira and Munro, 2011; Rosa-Ferreira et al., 2018). Strikingly, the number of Ilp2-GFP-marked DCVs in peripheral nerve motor axons and terminals of *Arl8*^*e00336*^ null larvae was severely reduced, with the remaining axonal DCVs being predominantly static (Fig. 7A, 3B-E, Video 6). The loss of axonal and synaptic DCVs was most likely due to a failure in the ability to exit the cell bodies, as Ilp2-GFP was still abundant in the VNCs of *Arl8* mutant animals (Fig. 7B). This mirrors the behavior of lysosome-related presynaptic carriers in *Arl8* nulls (Vukoja et al., 2018) and resembles the disruption of the DCV distribution in loss-of-function mutants for the Arl8-linking (Vukoja et al., 2018; Wu et al., 2013) unc-104 motor (Barkus et al., 2008; Lim et al., 2017; Pack-Chung et al., 2007). The few axonal DCVs that were still mobile exhibited strongly reduced run speeds, dramatically extended pause durations, and severely reduced processivity (Fig. 7A and Fig. 3D-E, Fig. S6A). Interestingly, despite the severity of the *Arl8*^*e00336*^ phenotype, we did not observe the relative reduction in retrograde DCV flux which characterized *Rab2*^*Δ1*^ mutants. In contrast to *Arl8*, axonal transport of DCVs in *Rab7*^*Δ1*^ null larvae was similar to wild type (Fig. 7A and 3B-E, Video 6). This further confirms that the observed DCV transport phenotypes in *Rab2* and *Arl8* mutants are not a secondary consequence of blocked fusion of LEs and autophagosomes with lysosomes.

**Figure 7.**
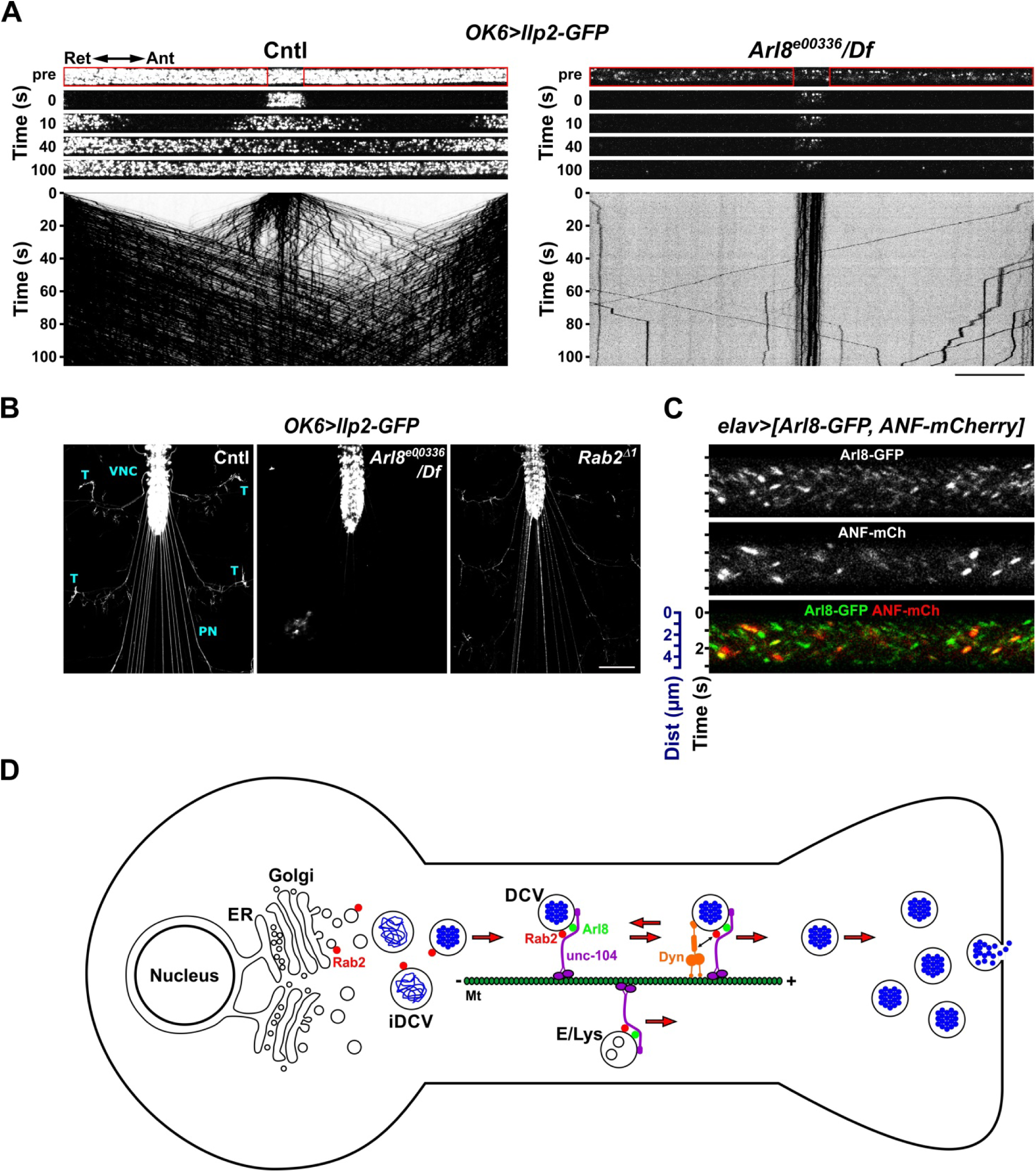
DCV axonal transport depends on Rab2 and Arl8. **(A-C)** Arl8 localizes to DCVs and is required for their axonal transport. **(A)** Transport of Ilp2-GFP-positive DCVs in motor axons of A7 nerves. DCVs were imaged before (Pre) and at the indicated time points after photobleaching the lateral areas (*red boxes*). The corresponding kymographs (inverted images) are shown below. See (Fig. 3) for quantification of the data in **(A). (B)** Sum projections of the anterior aspect of fillet-dissected larvae expressing Ilp2-GFP in motor neurons (live imaging). Note that in *Arl8* the Ilp2-GFP signal is mostly absent from peripheral nerves (PN) and terminals (T), but remains in the VNC. Images are representative of the following number of preparations: Cntl, 6; *Arl8*^*e00336*^*/Df*, 3; *Rab2*^*Δ1*^, 3. **(C)** Co-localization of Arl8-GFP with the DCV cargo marker ANF-mCherry on transported organelles in peripheral nerve axons, visualized as in Fig. 6, **(A and B). (D)** Proposed cellular functions of Rab2 and Arl8 in peptide-secreting neurons. Rab2 is present on TGN-associated vesicles and participates in the biogenesis and maturation of immature DCVs (iDCV). Arl8 is then recruited to DCVs and is responsible for their exit from the cell body and subsequent anterograde transport through the axon by recruiting the Kinesin-3/unc-104 motor (Vukoja et al., 2018; Wu et al., 2013). Rab2 also associates with DCVs and regulates their bidirectional transport through axonal shafts by interacting with unc-104 and possibly dynein and/or Kinesin-1/Khc (not shown). This process is important for maintaining a pool of DCVs at distal release sites. Arl8 (Farias et al., 2017; Rosa-Ferreira et al., 2018; Vukoja et al., 2018) and Rab2 are also required for the axonal transport of endosomal and lysosomal organelles (E/Lys). Scale bars: **(A)**, 20 µm; **(B)**, 200 µm. Dist, distance.

In light of Arl8’s critical role in axonal transport of DCVs, and the general assumption that Arl8 is primarily a lysosomal GTPase, we sought to establish its localization in relation to axonal DCVs. Imaging of peripheral nerve axons using the same method as that used to visualize fluorescently tagged Rab2 on moving organelles revealed neuronally expressed Arl8-GFP to be present on transiting ANF-mCherry-positive DCVs (Fig. 7C).

The fact that the late endosomal/lysosomal small GTPases Rab2 and Arl8 control axonal transport of both DCVs and lysosomes (Farias et al., 2017; Vukoja et al., 2018), as well as the close ultrastructural similarity of lysosome-related presynaptic carriers to classically described DCVs (Vukoja et al., 2018), raises the question whether these organelles are to some extent identical. Co-imaging of RFP-Spinster with Ilp2-GFP did indeed reveal some instances of transiting axonal DCVs clearly labeling with this lysosomal marker (Fig. S10A), but these events were rare. Likewise, brp-GFP and ANF-mCherry only coincided in a minor proportion of transport events involving active zone transport vesicles (Fig. S10B).

Collectively, these findings show that Arl8 is present on DCVs and is critically required for their axonal transport, similar to Arl8’s role in the transport of lysosomes and lysosome-related presynaptic carriers. However, DCVs are for the most part distinct from lysosomal organelles including those responsible for presynaptic biogenesis.

## DISCUSSION

DCVs are specialized organelles that underpin virtually all cellular signalling employing peptide hormones and neurotransmitters. This mode of signalling is especially prevalent in the nervous system, making highly critical the long-range motility of DCVs through axonal projections, from where most DCV exocytosis takes place (Persoon et al., 2018). However, the mechanisms controlling cytoskeletal transport of DCVs to their targets remain underexplored, especially regarding the upstream regulatory proteins that control molecular motors. Here we identify the small GTPases Rab2 and Arl8 as critical regulators of axonal DCV transport (Fig. 7d).

In addition to controlling DCV transport, we demonstrate that Rab2 is also required for efficient axonal transport of endo-lysosomal organelles. Likewise, Arl8 is a central regulator of lysosomal transport (Farias et al., 2017; Rosa-Ferreira and Munro, 2011; Vukoja et al., 2018). This illustrates an intriguing parallel between the biology of DCVs and lysosomes, reinforced by the fact that Rab2 is involved the biogenesis of both lysosomes (Lorincz et al., 2017; Lund et al., 2018) and DCVs (Edwards et al., 2009; Sumakovic et al., 2009; Topalidou et al., 2016). This and other similarities in the biogenesis of DCVs and lysosomes, such as the shared requirement for AP-3, Vps41 and BLOC-1 (Asensio et al., 2010; Asensio et al., 2013; Hao et al., 2015), may imply an evolutionary relationship between DCVs and lysosomal organelles. As Arl8 is also required for the biogenesis of lysosomes (Khatter et al., 2015; Lund et al., 2018), an interesting question for future studies would be if Arl8 plays a role in the formation or maturation of DCVs.

The picture emerging from our findings is that Rab2 exerts extensive control over DCV biology, sustaining both their formation and subsequent intracellular transport. The recruitment of Rab2 to maturing DCVs could conveniently position this GTPase for regulation of the transport motor complex. The fact that Rab2 and other transport-associated proteins such as unc-76 are targeted by the DIMM transcription factor (Hadzic et al., 2015) suggests that the cellular differentiation program initiated by DIMM to establish the high-capacity, far-projecting secretory phenotype in neurons (Hamanaka et al., 2010) includes upregulation of the axonal transport machinery to match the logistic challenge of supplying DCVs to distal release sites.

Importantly, our work also demonstrates that Arl8 and Rab2 have distinct functions in the orchestration of DCV transport. While Arl8 is critically required for driving DCV exit from somata and is a prerequisite for axonal motility, Rab2 supports bidirectional transport of circulating DCVs in the axonal shaft, especially in the retrograde direction. When driving lysosomal transport, Arl8 is known to recruit and activate Kinesin-3/unc-104 (Guardia et al., 2016; Vukoja et al., 2018; Wu et al., 2013), which is also the main anterograde DCV motor (Barkus et al., 2008; Lim et al., 2017; Lo et al., 2011). Consistent with this notion, the unc-104 loss-of-function DCV transport phenotype closely resembles that which we observe in Arl8 nulls (Barkus et al., 2008). Our results indicate that active Rab2 is likely involved in recruiting or activating unc-104 by participating in a molecular complex with this motor. However, the disproportionate effects of manipulating Rab2 on retrograde DCV transport also suggest that it may regulate DCV-related dynein function, consistent with previous work linking mammalian Rab2 to recruitment of dynein to ER-Golgi intermediates (Tisdale et al., 2009).

Dysfunction of human Rab2A has been linked to autism spectrum disorders (ASD) (Iossifov et al., 2012; Sanders et al., 2012; Takata et al., 2016), a set of high-heritability neurodevelopmental conditions associated with perturbations in the pro-reproductive and pro-social oxytocin (OT) and Arginine Vasopressin (AVP) neuropeptide systems (Meyer-Lindenberg et al., 2011). OT and AVP-producing neurons in the mammalian hypothalamus extend axonal projections to forebrain regions to regulate complex social behaviors. In our experiments, *Rab2* loss in flies depleted presynaptic DCV stores due to accumulation of stalled DCVs in the axon and reduced transport of active zone components to synaptic terminals. It is tempting to speculate that similar transport defects contribute to the ASD phenotype in individuals carrying deleterious *Rab2A* alleles.

In summary, our results uncover the mechanisms controlling axonal transport of DCVs, revealing surprising parallels between DCV and lysosomal motility.

## MATERIALS AND METHODS

### Fly husbandry, genetics, and stocks

Flies were reared on Nutri-Fly™ Bloomington Formulation medium (Genesee Scientific, San Diego, California) at 26°C, except for RNAi experiments, where they were reared at 29°C. In RNAi experiments, *UAS-Dicer-2* was co-expressed with the RNAi transgene. In *Rab2*^*Δ1*^ mutant experiments, larvae were reared on apple juice plates supplemented with yeast paste. For starvation experiments, L1 larvae aged 24-36 h after egg deposition were transferred to yeasted apple juice plates. After 48 h they were transferred to amino acid-poor starvation plates (agar, PBS, 1% (w/v) sucrose) (10-12 larvae/plate) and kept there for 24 h, before being dissected for immunohistochemistry. Fed control larvae were kept on yeasted apple juice plates. The full genotypes of animals used in experiments are listed in Table S1.

All DNA injections, selection of *P*-element and Phi31C transformants and selection of CRISPR/Cas9 knockouts (see below) was performed by BestGene Inc.. The 2xHA-TurboID-tagged transgenes encoding Rab2, Rab2^S20N^ and Rab2^Q65L^ were inserted into the M{3xP3-RFP.attP}ZH-86Fb attP site on the third chromosome using Phi31C transformation to ensure equal levels of expression.

The *Golgin-45*^*Δ1*^, *CG3703*^*Δ1*^ and *BicD*^*Δ1*^ knockout lines were generated using CRISPR/Cas9-catalyzed homology-directed repair according to the method outlined in (Gratz et al., 2014). In brief, two gRNA-expressing DNA constructs targeted near each end of the coding sequence of the genes were injected into *yw; nos-Cas9(II-attP40)* embryos expressing Cas9, together with a donor DNA construct based on the pHD-DsRed-attP plasmid. As a result, most of the coding sequence of the genes (X: 945355-942705 for *CG3703*, 3R: 9403771-9402693 for *CG9356*/*Golgin-45*, 2L: 17460549-17465046 for *BicD*) was replaced with a *DsRed* transgene. For *Golgin104* we used one gRNA-construct together with a matching homology-directed repair construct to target a DsRed transgene for insertion into the second coding exon of *Golgin104* between positions 3L:16350798-16350799, thus truncating the C-terminal 496 residues of the encoded protein, including the C-terminal Rab2-binding domain (Cattin-Ortola et al., 2017; Gillingham et al., 2014). Flies homozygous for the resulting *Golgin104*^*ins1*^ allele had reduced fecundity and were perceptibly smaller than wild type flies. The *BicD*^*Δ1*^ allele, in combination with the overlapping deficiency *Df(2L)Exel7068*, was hemizygously semi-lethal at the pupal stage, with 53 out of 199 tested pupae failing to eclose. We obtained similar results with hemizygotes over the same deficiency carrying the point nonsense-mutation allele *BicD*^*r5*^, previously described as amorphic (Ran et al., 1994). Flies homozygous for *Golgin-45*^*Δ1*^ or *CG3703*^*Δ1*^ did not display any obvious defects.

The *Rab2*^*Δ1*^ and *Vps39*^*Δ1*^ alleles and the *GFP-Rab2, HA-Rab2, UAS-pHlourin-Rab2* and *UAS-pHlourin-Rab2*^*Q65L*^ lines have been described (Lund et al., 2018), as have the *PICK*^*1*^ and *PICK*^*2*^ deletion alleles (Jansen et al., 2009) and the *UAS-GFP-Golgin245* line (Herlo et al., 2018). The *UAS-GFP-Rab7* (III) (42706), *UAS-GFP-Rab5* (III) (43336), *UAS-Rab11-GFP* (II) (8506), *UAS-mito-HA-GFP*.*AP/CyO* (8442), *386Y-Gal4* (25410), *Pin[Yt]/CyO; P{w[+mC]=UAS-mCD8::GFP*.*L}LL6* (5130), *P{w[+mW*.*hs]=GawB}elav[C155]* (458), *ccap-Gal4* (25685), *UAS-DCR2, w*^*1118*^ (24645), *Df(3R)Exel6178, P{w[+mC]=XP-U}Exel6178/TM6B, Tb*^*1*^ (7657), *OK6-Gal4* (64199), *BicD*^*r5*^, *cn*^*1*^*/CyO* (4553), *Df(2L)Exel7068/CyO* (7838), *lt*^*11*^*/In(2LR)Gla, wg*^*Gla*−*1*^ *PPO1*^*Bc*^ (26262), *Df(2L)lt45, PPO1*^*Bc*^*/SM1, lt*^*16*^ (26187), *PBac{RB}Arl8*^*e00336*^*/TM6B, Tb*^*1*^ (17846), *Df(3R)BSC222/TM6B, Tb*^*+*^ (9699), *Df(3L)BSC649/TM6C, Sb*^*1*^, *cu*^*1*^ (25739), *UAS-spin*.*myc-EGFP* (III) (39668), *UAS-brp-GFP* (III) (36292) lines were obtained from the Bloomington Stock Collection. The *UAS-Rab2-RNAi* line (v105358) was from the Vienna *Drosophila* Resource Center. The *Rab7*^*Δ1*^ line (Hegedus et al., 2016) was a gift from Dr. Gábor Juhász (Eötvös University, Budapest, Hungary), *UAS-mCherry-Rab7* a gift from Dr. Jui-Chou Hsu (National Tsing-Hua University, Hsinchu, Taiwan), UAS-spin-RFP (Johnson et al., 2015) a gift from Dr. Graeme W. Davis (University of California, San Francisco, CA), *UAS-Arl8-GFP* (Vukoja et al., 2018) a gift from Dr. Stephan J. Sigrist (Freie Universität Berlin, Berlin, Germany), and *c929-Gal4* (Hewes et al., 2003) a gift from Dr. Paul Taghert (Washington University, St. Louis, MO). The *Burs-Gal4* driver (Peabody et al., 2008) was a gift from Dr. Benjamin H. White (National Institute of Mental Health, Bethesda, USA), and the *Ilp2-Gal4* driver (Rulifson et al., 2002) a gift from Dr. P. Shen (University of Georgia, Athens, GA). The *UAS-ANF-EMD* (third chromosome insertion) (Rao et al., 2001) and *UAS-Ilp2-GFP* (second chromosome insertion) (Wong et al., 2012) lines were a gift from Dr. Edwin Levitan (University of Pittsburgh, Pittsburgh, PA).

### Molecular biology

To generate the RNAi-resistant UAS-*Rab2*^*RNAi-insens*^ transgenic construct, a *Rab2* coding region cassette was synthesized by GenScript (Nanjing, China) in which all degenerate codons were changed into synonymous codons by base pair substitution. This cassette was amplified and cloned into the pUAST vector. pUAST-pHluorin-Rab2^S20N^ was generated by GenScript by mutagenizing the pre-existing pUAST-pHluorin-Rab2 construct (Lund et al., 2018). To generate the *UAS*-*IA2-GFP* transgenic construct, we first PCR-amplified the EGFP coding region from the pEGFP-N2 plasmid and inserted it into the pUAST vector. We then amplified the IA2 coding region from cDNA clone GH05223 and inserted it into the resulting plasmid to obtain the final pUAST-IA2-GFP construct. The pUAST-ANF-mCherry construct was generated by GenScript by synthesizing the ANF-mCherry coding region and inserting it into the pUAST vector. Likewise, the pUASTattB-2xHA-TurboID-Rab2, pUASTattB-2xHA-TurboID-Rab2^S20N^ and pUASTattB-2xHA-TurboID-Rab2^Q65L^ constructs were generated by GenScript.

For CRISPR/Cas9-mediated mutagenesis, guide RNA (gRNA) target sequences (Golgin104/CG4925, GTAATCATTGCGTACCGCTT; Golgin-45/ CG9356 #1, GCAGGTGATTGCACGTAGTG; Golgin-45/CG9356 #2, GTTCTCGGGTCGTCTGTACA; CG3703 #1, GACAGCTACGTGGCCCGCAC; CG3703 #2, GACGCGAATAGCGCCAGTTC; BicD #1, AACTACTACCGAAAGTGCTC; BicD #2, AACCGCCCAAGAGCGCATAA) were inserted into the pBFv-U6.2 vector to obtain gRNA constructs targeting *CG3703, Golgin104, Golgin-45* and *BicD*. The donor DNA constructs for homology-directed repair were produced by inserting appropriate ∼1kb homology arms flanking the Cas9 cleavage sites into the pHD-DsRed-attP vector. All DNA work for CRISPR/Cas9 mediated mutagenesis was performed by GenScript.

### Immunohistochemistry

For immunohistochemistry preceding confocal imaging, L3 larvae or pharate adults were dissected in phosphate-buffered saline (PBS) containing 137 mM NaCl, 2.7 mM KCl, 1.5 mM KH_2_PO_4_, 6.5 mM NaH_2_PO_4_, pH 7.4. For experiments involving visualization of DCV cargos in IPCs, isolated dissected brains were briefly stored in Schneider’s insect cell medium (Life Technologies, A820) supplemented with 5% heat-inactivated fetal bovine serum (FBS) at room temperature (RT) prior to fixation. Whole animal fillets or isolated brains were then fixed in 3.7% formaldehyde in PBS at RT for 45 min, or on ice for 2 h, followed by 6×10 min washes in PBX (PBS with 0.3% Triton-X 100 [Sigma-Aldrich, T8787]) at RT. After a 2 h blocking step in blocking buffer (PBX with 5% normal goat serum, Sigma-Aldrich, G9023) at RT, fillets and brains were incubated 24-72 h with primary antibodies in blocking buffer at 4°C under gentle agitation. They were then washed 6 × 10 min in PBX, incubated with secondary antibodies in blocking buffer for 2 h at RT, and again washed 6 × 10 min in PBX. Finally, the specimens were washed 2 × 5 min in PBS and mounted in ProLong® Gold antifade reagent (Life Technologies, P36934). Antibodies used for immunohistochemistry and their working concentrations are listed in Table 1.

**Table 1.**
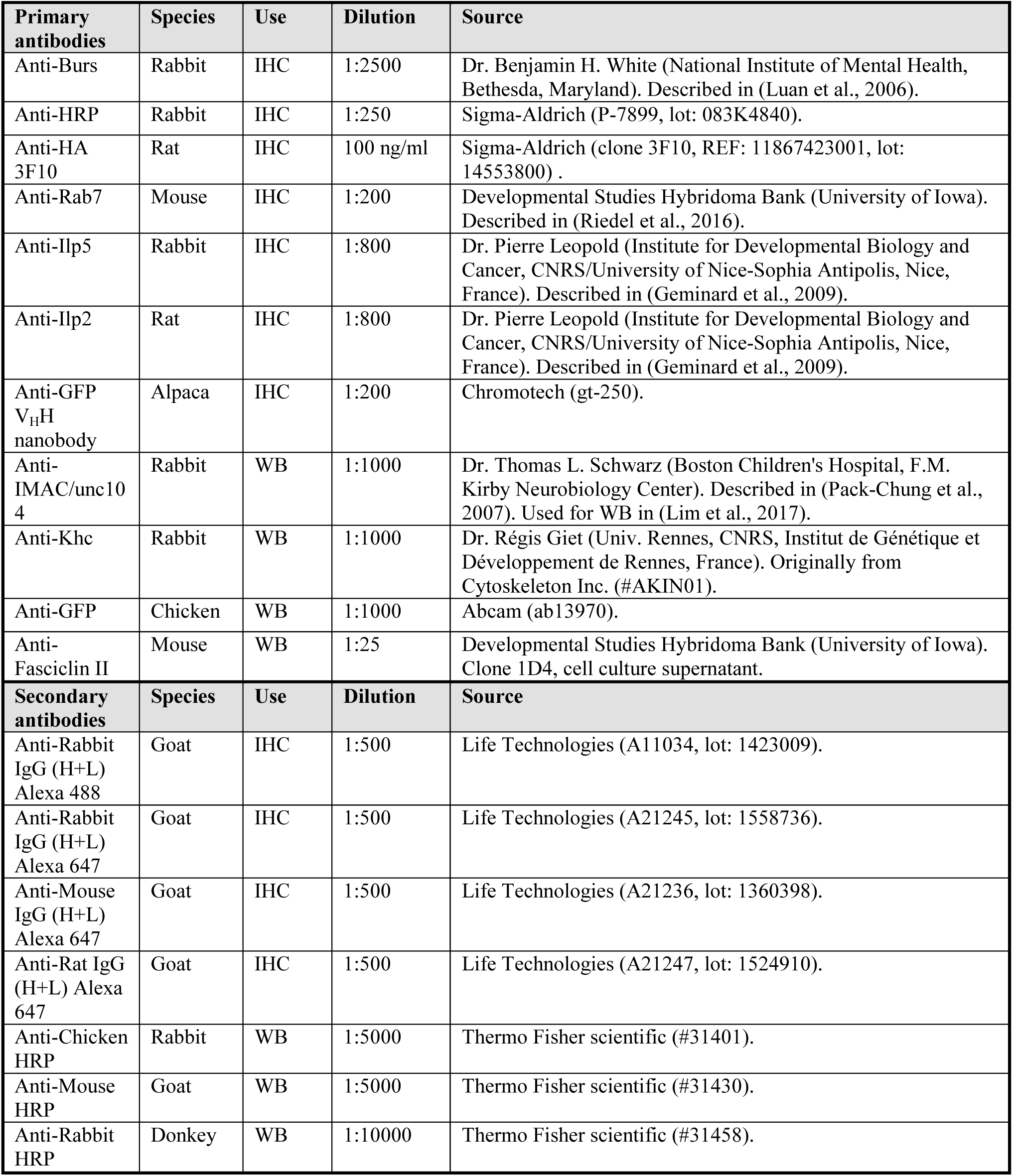
List of antibodies.

Fluorescent protein chimeras in fixed tissues were imaged directly without the use of antibodies.

### In vivo biotinylation and purification of biotinylated proteins

Flies expressing TurboID-Rab2, TurboID-Rab2^S20N^ or TurboID-Rab2^Q65L^ under control of the pan-neuronal *elav-Gal4* driver and *w*^*1118*^ control flies were reared on Nutri-Fly™ Bloomington Formulation medium supplemented with 100 µM Biotin. Adult flies were collected 0-3 days after eclosion, flash-frozen in liquid nitrogen and stored at -80°C.

For purification of biotinylated proteins, 1-2 ml of frozen flies were transferred to pre-cooled dounce homogenizers on ice, quickly dounced 5x, then dounced 15x in 3 ml RIPA lysis buffer (50 mM Tris (pH 7.5), 150 mM NaCl, 0.5% Sodiumdeoxycholate, 1.0% NP-40, 0.1% SDS, 1 mM DTT, cOmplete Protease Inhibitor Cocktail [1 tablet/25 ml](Roche, Ref: 11836145001), 1 mM PMSF). Lysates were then incubated for 30 min on ice and again dounced 15x. Hereafter the lysates were centrifuged 3 times for 15 min at 50.000g to remove insoluble debris. Protein concentrations in the resulting lysate supernatants were measured using the BCA assay (Pierce™ BCA Protein Assay Kit, Thermo Fisher Scientific, Ref: 23225, Lot: UE284353) and adjusted to 2.0 mg/ml. SDS was then added from a 10% stock solution to increase the SDS concentration to 0.5%. 1 ml of each lysate was pre-cleared for 1 h at 4°C under rotation with 100 µl Sepharose 4B beads, previously equilibrated in high-SDS RIPA buffer (50 mM Tris (pH 7.5), 150 mM NaCl, 0.5% Sodiumdeoxycholate, 1.0% NP-40, 0.4% SDS). Sepharose 4B beads were then removed by gentle centrifugation and supernatants were incubated over night at 4°C under rotation with 50 µl Dynabeads MyOne StreptavidinT1 (Thermo Fisher Scientific, Ref 65601, Lot: 00804134), previously equilibrated in high-SDS RIPA with 0.05% BSA. Dynabeads were then magnetically concentrated and washed 2x with 1 ml high-SDS RIPA buffer at RT, then transferred to fresh sample tubes in a third wash of high-SDS RIPA, washed 2x with SDS wash buffer (50 mM Tris pH 7.5, 2.0% SDS) and finally 2x with high-SDS RIPA buffer. Biotinylated proteins were eluted from the beads by incubation in elution buffer (50 mM Tris, 150 mM NaCl, 1.0% SDS, 1 mM biotin, 1 mM DTT) for 5 min at 95°C.

### Western Blotting

For western blotting of ANF-GFP, late L3 larval CNS with the ring gland and aorta attached were dissected out in PBS and stored in Schneider insect cell medium with 5% heat-inactivated FBS prior to freezing. Upon thawing, samples were immediately boiled in loading buffer, run on SDS-PAGE gels and electroblotted to PVDF membranes. Each sample consisted of pooled tissues from 5 individuals. Membranes were incubated in blocking solution consisting of 5% v/w Skim Milk Powder (Sigma-Aldrich #70166-500G) in PBS with 0.05% Tween 20 (PBST) for 1 h at RT and then probed with primary antibody at 4°C over night in blocking solution. They were then washed in PBST, probed with secondary antibody for 1 h at RT in blocking solution and again washed in PBST. For western blotting of isolated biotinylated proteins approximately half of the eluate from each binding reaction were boiled with SDS-PAGE loading buffer, run on SDS-PAGE gels and electroblotted to PVDF membranes, which were probed with antibodies as described above. Blots were developed using Amersham ECL Prime Western Blotting Detection Reagent (RPN2232) or SuperSignal ELISA Femto Maximum Sensitivity Substrate (Thermo Scientific, Ref: 37075, Lot: UO282555) and the resulting chemiluminescence imaged with an Alpha Innotech MultiImage Light Cabinet (Alpha Innotech, San Leandro, CA) equipped with a CCD camera. Antibodies used for western blotting and their working concentrations are listed in Table 1.

### Nanobody labeling for dSTORM

For dSTORM superresolution imaging of ANF-EMD-labeled DCVs in peripheral nerves, L3 larval fillets were prepared on Sylgard slabs in modified HL3 solution. The specimens were fixed in 3,7% formaldehyde in PBS for 45 min at RT, followed by two quick rinses and one longer 5 min incubation in quenching solution (PBS with 20 mM Glycine and 50 mM NH_4_Cl) to quench background fluorescence. Specimens were washed 7 × 10 min in PBX, incubated in blocking solution consisting of PBX with 5% Donkey Serum and 1% BSA for 3 h at RT and then incubated with an anti-GFP V_H_H single domain antibody/nanobody (Chromotech, gt-250) custom-conjugated to Alexa 647 (1:200) in blocking solution for ∼72 h at 4°C. They were then washed 10 × 10 min in PBX and fixed again in 3,7% formaldehyde in PBS for 45 min at RT, followed by two rinses and two longer 5 min incubations in quenching solution. After a brief wash in PBS the fillets were stored in PBS at 4°C until imaging.

### dSTORM imaging and post processing

Larval fillets were placed on a 17 nm thick poly-D-lysine coated coverslip and held in place with a metal weight. dSTORM images were taken on an ECLIPSE Ti-E epifluorescence/TIRF microscope (NIKON, Japan) utilizing excitation lasers at wavelengths 405 and 647 nm. The light was reflected by a dichroic mirror (z405/488/561/647 rpc) and emitted light filtered by a 710/80 nm bandpass filter (NIKON). Images were recorded with an EM-CCD camera (iXon3 897, Andor, United Kingdom). The objective used was a CFI Plan Apochromat VC 100X oil immersion objective (NIKON). dSTORM images were captured in imaging buffer containing β-mercaptoehtanol, cyclooctatetraene, and an enzymatic oxygen scavenger system (10% (w/v) glucose, 1% (v/v) beta-mercaptoethanol, 2 mM cyclooctatetraene, 50 mM Tris-HCl (pH 8), 10 mM NaCl, 34 μg/ml catalase, 28 μg/ml glucose oxidase). Videos of the fluorophore blinking were taken for a total of 20000 frames at 16 ms exposure time per frame.

Localizations were fit using the program ThunderSTORM (Ovesny et al., 2014), where localizations were detected with a threshold of 1.5 * standard deviation (Wave.F1) using the Local Maximum method. Drift correction was then performed with cross correlation. Images were generated by rendering the normalized Gaussian image with a 30x magnification through ThunderSTORM.

### Confocal microscopy

All confocal microscopy was carried out at the Core Facility for Integrated Microscopy (Department of Biomedical Sciences, University of Copenhagen) using LSM 700 and LSM 710 confocal microscopes (Carl Zeiss Microscopy GmbH, Jena, Germany) and the following objectives: Plan-APOCHROMAT 63x/1.4 Oil DIC, W APOCHROMAT 40x/1.0 DIC VIS-IR #421462-9900 Water dipping, Plan APOCHROMAT 20x/0.8 and EC Plan-NEOFLUAR 10x/0.3. Bleaching was performed using a 405 nm diode laser.

### Live imaging of axonal transport, DCV capture, and neuropeptide release

For live time lapse confocal imaging of axonal transport and synaptic capture of fluorescently labeled organelles, fillet dissections of L3 larvae or pharate adults were performed in modified HL3 solution ([HL3-EGTA] 70 mM NaCl, 5 mM KCl, 10 mM NaHCO_3_, 20 mM MgCl_2_, 5 mM trehalose (Merck KGaA, 108216), 115 mM sucrose (Sigma-Aldrich, S1888), 0.5 mM EGTA, 5 mM HEPES, pH 7.2. VNCs, peripheral nerves and terminals were then imaged directly in a Sylgard-coated dissection dish using the 40x water dipping objective. All live imaging was performed in HL3-EGTA unless otherwise indicated.

To examine DCV transport in peripheral nerves of larvae expressing *OK6*>Ilp2-GFP we chose a proximal region on the A7 nerve ∼0.5-1.0 mm from its exit from the VNC (or, in larvae expressing *386Y*>ANF-GFP, on the A8/9 nerve ∼0.5 mm from its exit). If required, the dissection dish was elevated on one side with Parafilm strips to orient the nerves horizontally. After fitting a 130 × 5.6 µm acquisition window (56 × 15 µm for *386Y*>ANF-GFP) to the nerve, all fluorescent signal except a central 10 µm nerve segment was photobleached once and time series were recorded with a frame rate of ∼5 frames per second (fps). A similar approach was used to examine the axonal transport of mito-GFP, GFP-Rab5, GFP-Rab7, spin-GFP and brp-GFP, though no bleaching was used in the case of brp-GFP and framerates were set depending on the quality of the fluorescent signal of the marker. For FRAP experiments in *386Y*>*ANF-GFP* larvae, an 18 µm long segment of the A8/9 nerve was bleached and recovery monitored in a 90 µm long acquisition window every 10 s.

For visualization of small amounts of GFP-Rab2, pHlourin-Rab2 or Arl8-GFP on organelles being transported in peripheral axons, a ∼130 µm long region of A7 or A6 peripheral nerves was photobleached and then imaged with the scanning direction parallel with the nerve and with a pixel dwell of 2.15-2.70 µs, an averaging number of 16 and a pixel size of 0.008 µm^2^. When co-imaging with ANF-mCherry, only the central ∼25-30 µm of the bleached region was used as the post-bleaching acquisition window. Large amounts of ANF-mCherry accumulated in extensive tubulo-vesicular organelles in glial sheath cells in the larval peripheral nerves when expressed in neurons using the *elav*-Gal4 driver, likely due to endocytic uptake of DCV cargo exocytosed from the axons. These organelles were highly immobile and the mCherry signal contained in them was bleached away as part of the experimental procedure, thus not interfering with the visualization of transiting DCVs in the axons contained inside the nerve.

For flux measurements of Ilp2-GFP-labeled DCVs in the distal axon supplying muscle 3 type Ib terminals, confocal time series of the terminal nerve branch in segment A4 were recorded with a frame rate of 3.2 fps for 500 frames. Simultaneously, the nerve branch was photobleached every 25 frames using a circular bleach box 10 µm in diameter. Ilp2-GFP-positive vesicles entering the area covered by the bleach box from either side between each bleach cycle were counted, while the bleaching eliminated vesicles from previous imaging cycles. This method allowed for a more accurate counting than simple time series, due to the high density of Ilp2-GFP-labeled DCVs in this axon. For flux measurements of *386Y*>ANF-GFP-labeled DCVs, also in the distal axon supplying muscle 3 type Ib terminals, simple confocal times series were recorded with a frame rate of 1.0 fps for 500 frames, as the observed flux of labeled vesicles was lower than for *OK6*>Ilp2-GFP.

For flux measurements of Ilp2-GFP-labeled DCVs in proximal axons of pharate adult stage Bursicon-secreting neurons, we recorded confocal time series of posterior lateral nerve trunks near their exit from the VNC with a frame rate of 8.3 fps for a total of 2000 frames.

For detection of co-transport of Ilp2-GFP and spin-RFP or ANF-mCherry and brp-GFP in axonal organelles we chose intersegmental nerve branches in fillet dissected OK6>Ilp2-GFP, spin-RFP or elav>ANF-mCherry, brp-GFP larvae, respectively. A 40 µm long stretch of nerve was photobleached to remove the fluorescence from static structures that might obscure transiting organelles and then imaged at 2.9 fps (Ilp2-GFP and spin-RFP) or 1.2 fps (ANF-mCherry and brp-GFP).

To examine DCV dynamics in NMJs, muscle 3 Ib terminals in segments A3-5 og filleted L3 larvae expressing *OK6*>Ilp2-GFP were imaged employing a wide optical slice (2.1 µm, corresponding to 1.85 Airy units) and a frame rate of 1.1 fps. To measure capture, the entire terminal was photobleached, followed by a pause of a duration manually adjusted such that 2-9 vesicles were allowed to enter the terminal from the distal axon. The terminal was then imaged (800 frames) while photobleaching the distal axon every 20th frame to bleach any new vesicles entering the terminal. The duration of each bleach cycle was 1-1.5 s depending on the shape of the bleach box. To measure the efflux of vesicles from the most distal bouton, the entire terminal and distal axon except for the distal bouton was photobleached, followed by time-lapse microscopy at settings identical to those described above, while photobleaching the distal axon every 20 frames.

For neuropeptide release experiments, L3 larvae expressing *OK6*>Ilp2-GFP were dissected in Ca^2+^-free saline (128 mM NaCl, 2 mM KCl, 4 mM MgCl_2_, 35.5 mM sucrose, 0.5 mM EGTA and 5 mM Na HEPES, pH 7.2) and Ilp2-GFP fluorescence was then recorded in Ib terminals of muscle 3 NMJs in segments A3-5 as fast confocal stacks. Release was evoked by depolarization obtained by replacing the bath solution with high-K^+^ saline (130 mM KCl, 1.8 mM CaCl_2_, 4 mM MgCl_2_, 35.5 mM sucrose and 5 mM Na HEPES, pH 7.2) for 3 min, followed by replacement of bath solution with Ca^2+^-free saline to terminate the stimulus and allow for muscle relaxation. Confocal stacks were recorded immediately before stimulation and immediately after the washout of the high-K^+^ saline (0 min), as well as at 5, 10 and 15 minutes after termination of stimulation.

### Image processing and analysis

*FRAP experiments*. To quantify the FRAP of bleached peripheral nerves in larvae expressing *386Y*>ANF-GFP we measured the fluorescence intensity in the bleached area and performed double normalization of the raw intensity (*I*_*raw*_) to the pre-bleach intensity (*I*_*pre*_) and to two non-bleached segments of nerve (*I*_*cont*_) to control for bleaching during acquisition, while subtracting the baseline intensity (*I*_*base*_) defined as the intensity of the bleached area immediately after bleaching (*t* = 0):

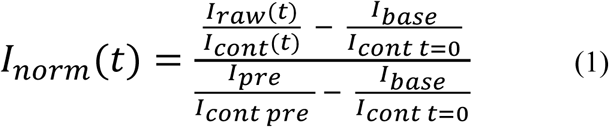

A double exponential decay equation was fit to the data using the ImageJ curve fitting tool.

### Mid-nerve axonal transport

Trajectories of trafficking fluorescent organelles entering the bleached areas were traced manually on kymographs by drawing them as segmented lines in Fiji/ImageJ (Schindelin et al., 2012). Since the complexity caused by the many trafficking organelles generally precluded complete tracing of individual trajectories, all runs or pauses were pooled for each time series. Segmented line coordinates were extracted and imported into Excel (Microsoft) for further analysis. Segments were counted as pauses when organelle speeds in either direction did not exceed 0.015 µm/s (ANF-GFP and Ilp2-GFP) or 0.0075 µm/s (mito-GFP, GFP-Rab5, GFP-Rab7, and spin-GFP).

### Quantification of static and dynamic organelles

To count the number of static vesicles in the central unbleached area (CUA) in mid-nerve experiments involving bilateral photobleaching, a separate kymograph was produced for each y-coordinate. As the frame height was 56 pixels, this resulted in a set of 56 kymographs per time series (a “kymostack”). Kymostacks made it easier to distinguish individual trajectories, since information about the organelle position along the y-axis was retained. Organelles were counted as static if they did not deviate more than 15 pixels from their starting point during the entire time series. This criterion was also used in the experiment involving brp-GFP. Dynamic organelles leaving the CUA were counted in standard kymographs as the number of trajectories crossing a vertical line one µm to the left (retrograde transport) or to the right (anterograde transport) of the CUA border. Alternatively, to obtain a sufficient number of trajectories for analysis, we counted organelles invading the bleached areas from the unbleached regions flanking the imaged field as the number of crossings at x-positions 400 pixels away from the left (anterograde transport) or the right (retrograde transport) edge.

### Visibility-enhancing image processing

Since the mCherry-Rab7 signal (Video 2) suffered heavily from bleaching during imaging of axonal transport, the contrast of each separate frame was adjusted by saturating 0.35% of the pixels before converting the series to a movie. The kymographs from time lapse imaging of *OK6*>Rab5-GFP, *OK6*>Rab7-GFP, and *OK6*>spin-GFP transport (Fig. 5) exhibited a steep bilateral intensity gradient, causing excessive saturation when contrasting the images to visualize the faint trajectories in the center of the images. Structures larger than 100 pixels were therefore filtered using the FFT bandpass Plugin in Fiji before producing the Figure. The red channel (spin-RFP) in the *OK6*>[Ilp2-GFP, spin-GFP] kymograph, and both channels in the *elav*>[brp-GFP, ANF-mCherry] kymograph, were bandpass filtered (3-100 pixels) to similarly compensate for uneven intensity.

### Synaptic capture and size estimation of vesicle pools

To quantify capture of DCVs following photobleaching of all synaptic boutons, the time points at which individual vesicles shifted between synaptic boutons were manually tabulated frame by frame, from which residence times were calculated. For experiments where all but the distal-most bouton were bleached, the total pool of vesicles in that bouton was estimated by dividing its pre-bleach intensity with the average intensity of nearby single vesicles after their exit from the bouton (number of vesicles used: *w*^*1118*^, 6.14 ± 0.637; *Rab2*^*Δ1*^ 5.16 ± 0.443; mean ± s.e.m.). Exponential fitting of the cumulated number of exiting vesicles was done with the Solver tool (Excel).

### Intensity measurements

Levels of Ilp2-ILP2, ANF-GFP, and the PICK1 or Futsch immunosignal were measured in confocal sum stacks. In some cases, thresholding following by particle analysis was applied within the region of interest to more precisely delineate the signal. Background subtraction was performed by measuring background intensity in a separate region or by employing the rolling ball subtraction plugin in Fiji (radius 50 pixels).

### Statistical analysis

ANOVA followed by Tukey’s honest significant difference test or Dunnett’s test was performed using JMP (SAS Institute Inc., Cary, NC). Student’s *t*-tests were exclusively two-sided and performed in Excel, assuming either equal or unequal variances (Welch’s test) as estimated with a preceding F-test. No data were rejected as outliers based on automated procedures, as apparent “outliers” often reflected a lognormal distribution. When required, data were logarithmically or square root transformed before statistical testing to obtain normal distributions with similar variances, verified by inspecting residuals. The level of statistical significance was set to 5%. To avoid temporal bias, care was taken to test different genotypes in mixed order. Based on experience with similar work, sample sizes were generally kept at five or more in order to ensure sufficient statistical power.

## DATA AVAILABILITY

Data supporting the findings of this manuscript are available from the corresponding author upon reasonable request. The quantitative data underlying Figures and Supplementary Figures are provided as a Source Data file.

## ACKNOWLEDGEMENTS

We thank the staff at the Core Facility for Integrated Microscopy and Anders Bohl Pedersen for excellent technical assistance, and Kenneth Lindegaard Madsen for useful discussions. We thank the Vienna Drosophila Resource Center, the Bloomington Stock Center, and the following colleagues for fly lines and reagents: Gábor Juhász, Stephan J. Sigrist, Thomas L. Schwarz, Pierre Leopold, Graeme W. Davis, Benjamin H. White, Edwin Levitan, Suzanne Eaton, Jui-Chou Hsu, Paul Taghert, P. Shen and Régis Giet. We acknowledge the respective communities for maintaining the Fiji and Inkscape software. Supported by Beckett Fonden, Dagmar Marshalls Fond, Else og Mogens Wedell Wedellsborgs Fond, Læge Sofus Carl Emil Friis og Hustru Olga Doris Friis Legat, Novo Nordisk Fonden, Torben og Alice Frimodts Fond, and Aase og Ejnar Danielsens Fond.

## AUTHOR CONTRIBUTIONS

V.K.L. conceived and planned the experiments supervised by O.K.. V.K.L. performed experiments, assisted by R.C.A. (western blots) and O.K. (release experiments). M.D.L. contributed with dSTORM microscopy. O.K. and V.K.L. analyzed the data assisted by A.S. (ANF-GFP trafficking) and produced figures. V.K.L. and O.K. wrote the manuscript with input from all authors. O.K. and U.G. provided funding.

## VIDEO LEGENDS

**Video 1**. Rab2 loss disrupts dense core vesicle axonal transport. Live confocal imaging illustrating the disrupted transport of *OK6*>Ilp2-GFP-positive DCVs in motor neuron axons of *Rab2*^*Δ1*^ larvae and rescue of the transport by expression of a *Rab2* transgene. The lateral regions of the imaged segment of the A7 peripheral nerve were photobleached after the first frame as in (Fig. 3A) to enhance the visibility of DCVs exiting from the central unbleached region and entering from the sides. Ant, anterograde (towards synaptic terminals); Ret, retrograde (towards neuronal cell bodies). The Cntl and *Rab2*^*Δ1*^ experiments are identical to those shown in (Fig. 3A).

**Video 2**. Alterations in DCV axonal transport in *Rab2* nulls are not caused by rerouting of DCV cargo to late endosomes. Live confocal imaging illustrating the relation of *386Y*-Gal4-driven ANF-GFP (*green*) and mCherry-Rab7 (*red*) in peptidergic axons of wild type control (*top*) and *Rab2* null (*bottom*) larvae. The lateral regions of the imaged segment of the A8/9 peripheral nerve were photobleached after the first frame (*red boxes*) to enhance the visibility of moving organelles. Photobleaching through imaging was compensated for by applying contrast enhancement on each image (see Methods). Note the increase in static ANF-GFP-positive DCVs in the central unbleached area in *Rab2*^*Δ1*^ compared to controls. Note also the relatively low level of overlap between ANF-GFP and mCherry-Rab7-positive organelles in both control and *Rab2*^*Δ1*^ axons.

**Video 3**. Rab2 loss disrupts early endosome axonal transport. Live confocal imaging illustrating the disrupted transport of *OK6*>GFP-Rab5-positive EEs in *Rab2* null motor neuron axons. The lateral regions of the imaged segment of the A7 peripheral nerve were photobleached after the first frame as in (Fig. 3A) to enhance the visibility of moving organelles. Same experiments as shown in (Fig. 5A, *left*). Ant, anterograde (towards synaptic terminals); Ret, retrograde (towards neuronal cell bodies).

**Video 4**. Rab2 loss disrupts late endosome axonal transport. Live confocal imaging illustrating the reduced transport of *OK6*>GFP-Rab7-positive LEs in *Rab2* null motor neuron axons. The lateral regions of the imaged segment of the A7 peripheral nerve were photobleached after the first frame as in (Fig. 3A) to enhance the visibility of moving organelles. Same experiments as shown in (Fig. 5B, *left*). Ant, anterograde (towards synaptic terminals); Ret, retrograde (towards neuronal cell bodies).

**Video 5**. Rab2 loss disrupts axonal transport of lysosomes and lysosome-related vesicles. Live confocal imaging illustrating the reduced transport of *OK6*>Spinster-GFP-positive lysosomes and lysosome-related vesicles in *Rab2* null motor neuron axons. The lateral regions of the imaged segment of the A7 peripheral nerve were photobleached after the first frame as in (Fig. 3A) to enhance the visibility of moving organelles. Same experiments as shown in (Fig. 5C, *left*). Ant, anterograde (towards synaptic terminals); Ret, retrograde (towards neuronal cell bodies).

**Video 6**. Arl8, but not Rab7, is critical for axonal transport of DCVs. Live confocal imaging illustrating the disrupted transport of *OK6*>Ilp2-GFP-positive DCVs in motor neuron axons of *Arl8* null, but not *Rab7* null larvae. The lateral regions of the imaged segment of the A7 peripheral nerve were photobleached after the first frame as in (Fig. 3A) to enhance the visibility of moving organelles. Ant, anterograde (towards synaptic terminals); Ret, retrograde (towards neuronal cell bodies).

