## Supplemental figures and table for "Rab2 drives axonal transport of dense core vesicles and lysosomal organelles"

**Lund et al.**

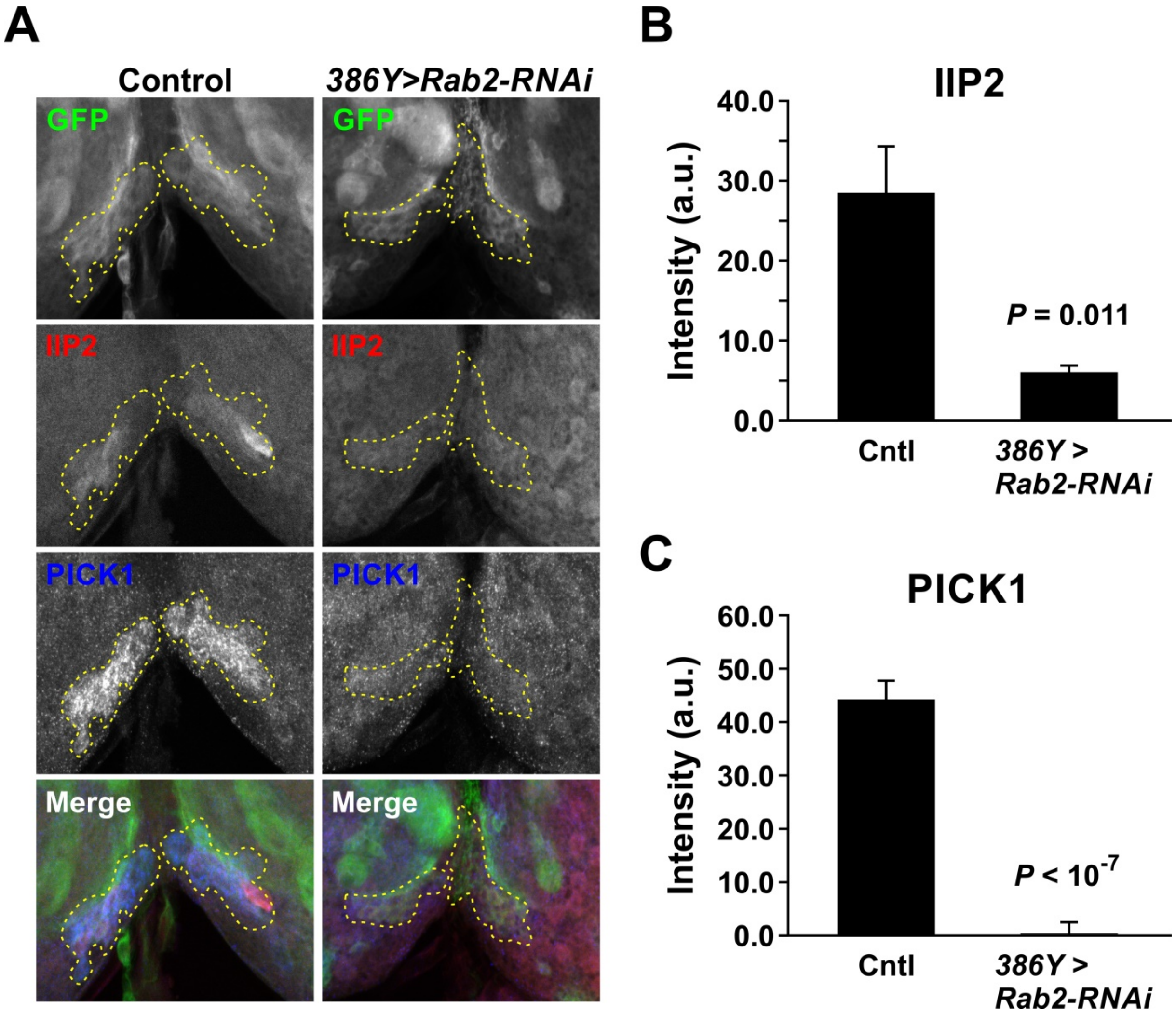


**Figure S1. Depletion of Rab2 in peptidergic neurons leads to a loss of PICK1 and Ilp2 from IPC somata**. **(A)** Peptidergic neurons in larval brains labeled with membrane-bound mCD8-GFP (*top*). *Left*, control; *right*, *Rab2* knockdown. IPC somata in pars intercerebralis are outlined (*dashed yellow lines*). *Second and third rows*, Ilp2 and PICK1 immunosignals, respectively. Anterior is downwards. Single optical slices. **(B and C)** Intensity of the Ilp2 **(B)** and PICK1 **(C)** immunosignals in the IPC somata in **(A)**. Data are mean ± s.e.m. from nine controls and seven *Rab2* knockdown preparations. Student's *t*-tests. A.u., arbitrary units. Scale bar: 20 µm.

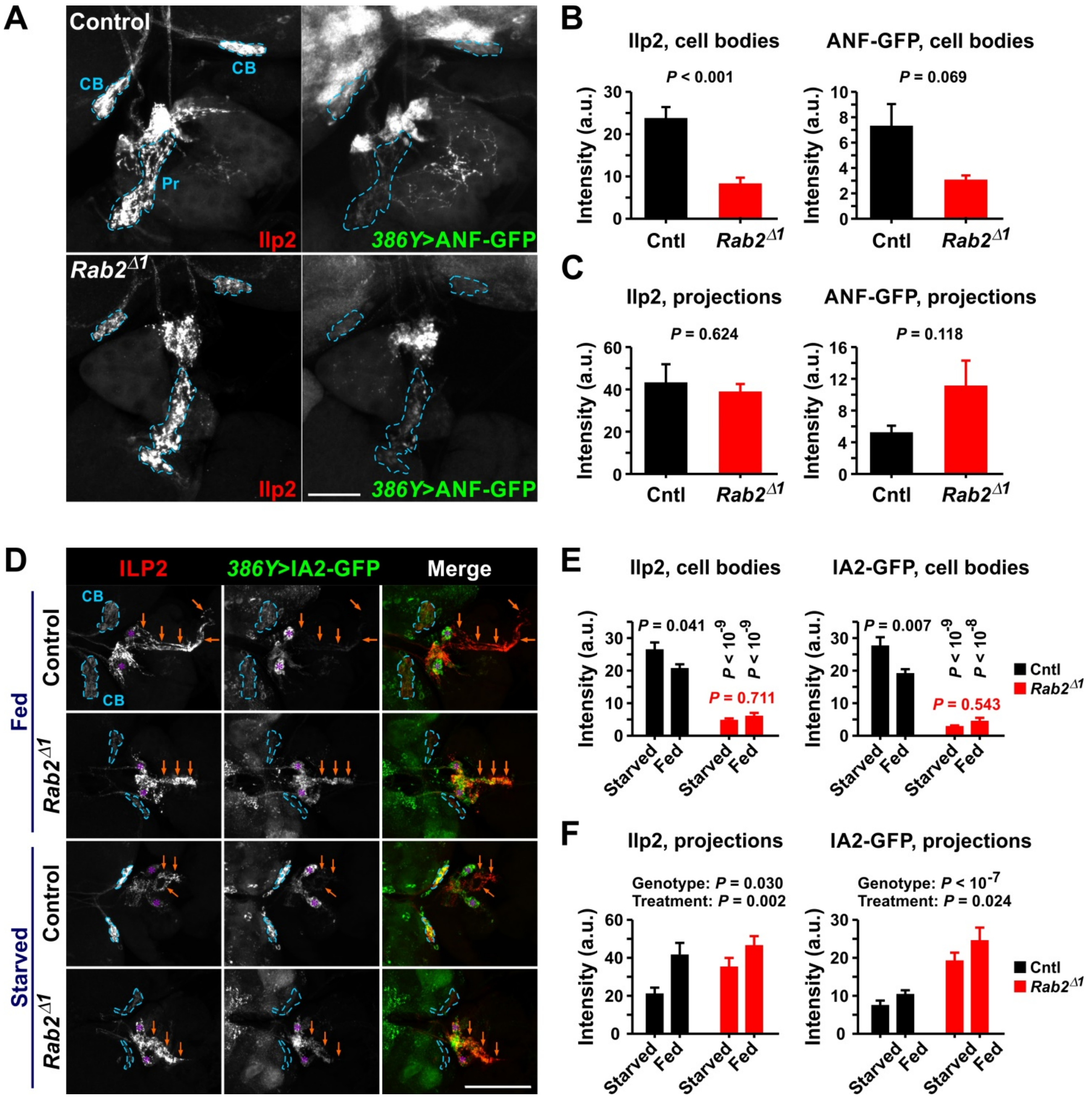


**Figure S2. Effects of Rab2 loss on DCV cargo in brain IPCs. (A)** Ilp2 immunoreactivity and *386Y*>ANF-GFP signal in cell bodies (CB) and axonal projections (Pr) of control (*top*) and $Rab2^{\Delta 1}$ larvae (*bottom*). **(B and C)** Quantification of signal intensities in **(A)**. Data (mean ± s.e.m.) from five control and six $Rab2^{\Delta 1}$ larval CNS preparations. Student's *t*-tests. **(D)** Ilp2 immunoreactivity and *386Y*>IA2-GFP signal in cell bodies (*dashed blue lines*) and projections (*blue arrows*) of brain IPCs from starved and fed control and $Rab2^{\Delta 1}$ mutant larvae. Corpora cardiaca marked by *blue asterisks*. **(E and F)** Quantification (mean ± s.e.m.) of signal intensities in **(D)**. Two-factor ANOVA, followed in **(E)** by Tukey's test. Each of the four combinations of genotype and feeding state are represented by seven to nine CNS preparations. Specific sample sizes are available from the provided Data Source file. A.u., arbitrary units. Scale bars: **(A)**, 40 µm; **(D)**, 100 µm.

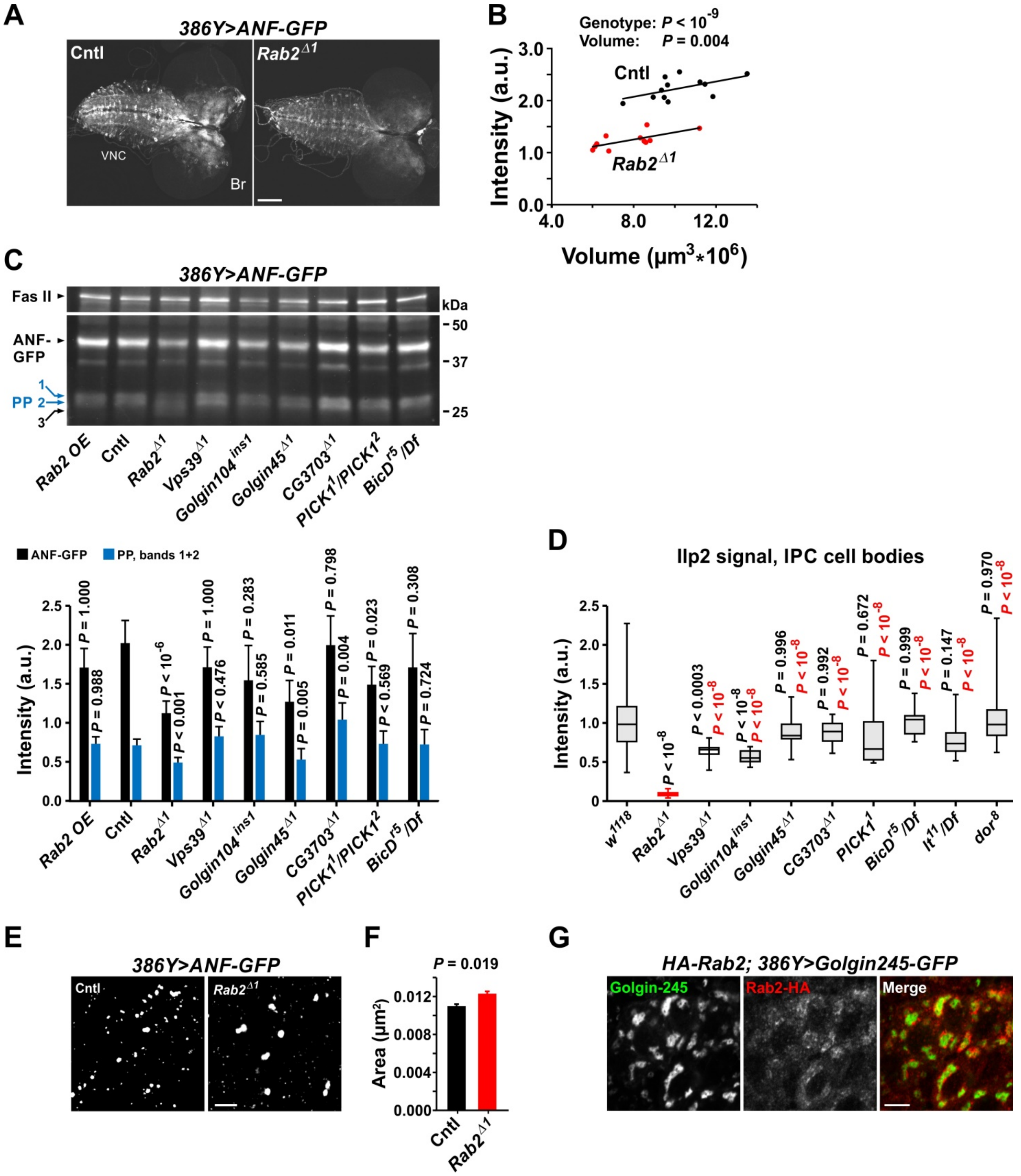


**Figure S3. Reduced levels and aberrant processing of neuropeptides, and perturbed DCV size distribution, in *Rab2*$^{\Delta 1}$ larvae.** **(A)** GFP in fixed brain (Br) and ventral nerve cord (VNC) of control and *Rab2* null mutant L3 larvae expressing ANF-GFP in peptidergic neurons. **(B)** Quantification of **(A)**. ANF-GFP intensity versus CNS volume. Data from 12 control and 11 *Rab2*$^{\Delta 1}$ CNS preparations. Two-factor ANOVA. **(C)** (*top*) Western blot of larval CNS from the indicated genotypes expressing ANF-GFP driven by *386Y*-Gal4. Fasciclin II used as loading control. The ~41 kDa band corresponds to ANF-GFP cleaved by an unknown neuropeptide-processing enzyme, whereas the cluster of bands at around ~26 kDa represents another set of ANF-GFP proteolytic products (PP 1, 2 and 3) {Rao, 2001 #418}. Note in the *Rab2*$^{\Delta 1}$ sample the marked increase in PP 3, and the decreased intensity of ANF-GFP and PP 1 and 2. Rab2 OE: *UAS-Rab2* transgene overexpressed in peptidergic neurons using the *386Y*-Gal4 driver that also controlled the ANF-GFP expression. **(C)** (*bottom*) Quantification of **(C)** (*top*), showing the intensity of the immunosignal for each genotype. Data are from six western blots, with each genotype included on at least three separate blots. Exact sample sizes are available from the Data Source file. Mean ± s.e.m. ANOVA followed by Tukey's test. *$P$<0.05: **$P$<0.005; ***$P$<0.0001; N.s., not significant. A.u., arbitrary units. **(D)** Box plot (min, lower quartile, median, upper quartile, maximum) of the Ilp2 intensity in IPC somata from starved larvae of the indicated genotypes, normalized to the *w*$^{1118}$ mean (see Table S1 for detailed genotypes). ANOVA followed by Tukey's test. *P*-values in *black* and *red* from comparisons with *w*$^{1118}$ and *Rab2*$^{\Delta 1}$, respectively. Each genotype represented by at least 9 CNS preparations. Exact sample sizes are available from the Data Source file. **(E)** Representative dSTORM images of peripheral nerves in control and *Rab2*$^{\Delta 1}$ mutant larvae expressing ANF-GFP driven by *386Y*-Gal4. **(F)** Quantification of **(E)**. Each genotype represented by three larvae, with 7-8 images analyzed per larva. Exact sample sizes are available from the Data Source file. Nested ANOVA. **(G)** HA-Rab2 tightly envelops TGN structures. Dorsal VNC from *HA-Rab2* pharate adult, immunostained for HA, and expressing the TGN marker Golgin245-GFP in peptidergic neurons. Scale bars: **(A)**, 100 µm; **(E)**, 1 µm; **(G)**, 5 µm.

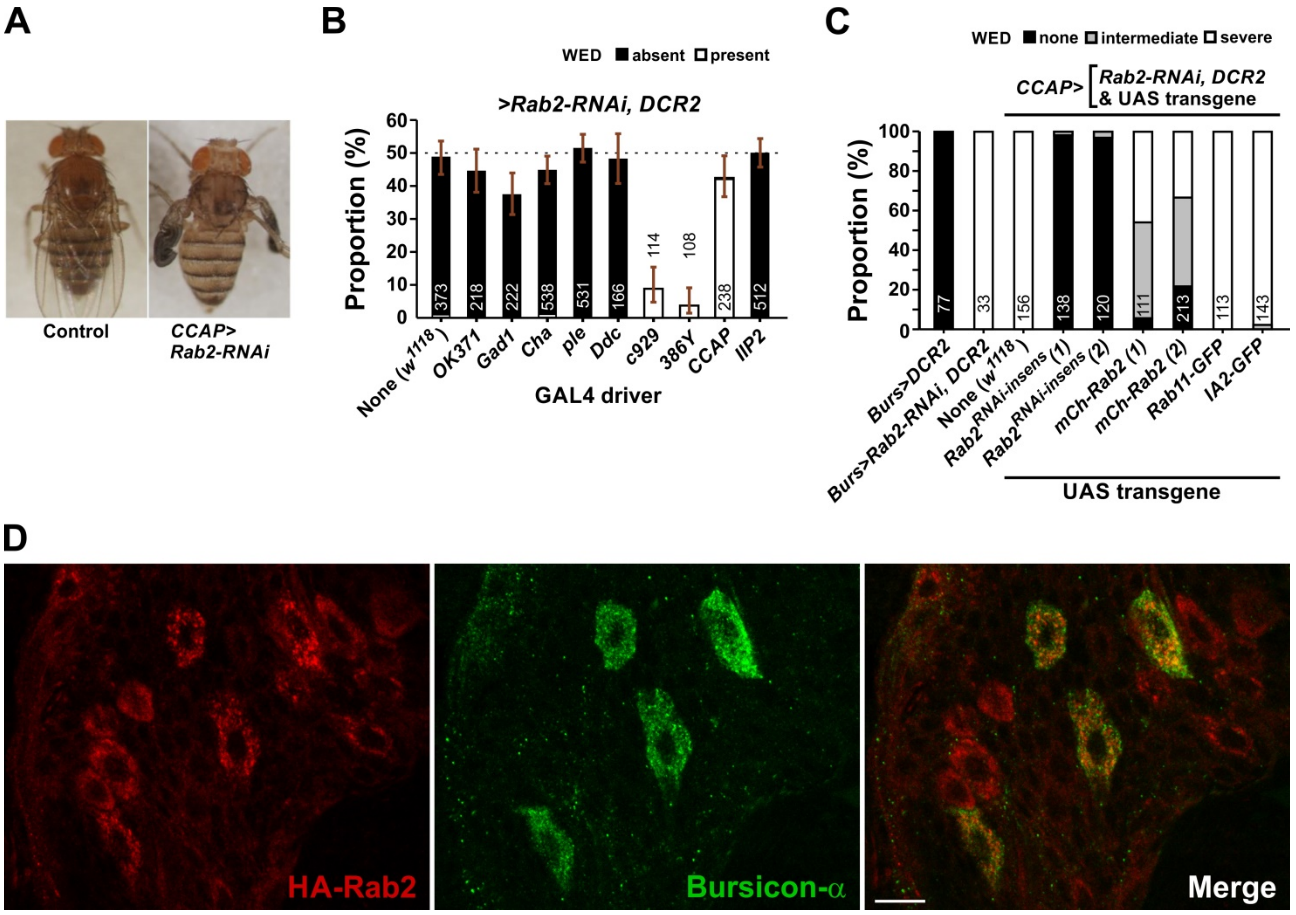


**Figure S4. Depletion of Rab2 in peptidergic neurons impedes viability and wing expansion in adult flies. (A)** Unlike in control flies (*left*), *Rab2* knockdown in neurons expressing crustacean cardioactive peptide (CCAP) as well as Bursicon blocked post-eclosion wing expansion (*right*). **(B)** Eclosion proportion (mean and 95% confidence interval) and degree of wing expansion defect (WED) after *Rab2* knockdown in neuronal populations employing specific neurotransmitters, driven by *OK371*-Gal4 (glutamate), *Gad1*-Gal4 (GABA), *Cha*-Gal4 (acetylcholine), *ple*-Gal4 (dopamine), *Ddc*-Gal4 (dopamine, serotonin). *386Y*-Gal4 (general peptidergic), *CCAP*-Gal4 (CCAP), or *Ilp2*-Gal4 (insulin-like peptide 2). A control group omitting a Gal4 driver is also included. Shown is the proportion of the total number of offspring (indicated *in or close to bars*) that carried the *Rab2* RNAi transgene rather than a balancer chromosome (see Methods). Without detrimental knockdown effects the expected eclosion proportion was 50%. **(C)** *Rab2* knockdown in a subset of CCAP neurons driven by *Burs*-Gal4 induces a severe wing expansion defect (WED), partially rescued by expressing two different *mCherry-Rab2* transgenes, and fully rescued by two different RNAi-insensitive *Rab2* transgenes. No rescue occurred when expressing Rab11-GFP or the IA2 DCV membrane protein. Severe wing expansion defect implies a complete absence of wing expansion, as in **(A)** (*right*). The number of flies evaluated is indicated *inside bars* for each genotype. **(D)** Dorsal surface of *HA-Rab2* pharate adult VNC immunostained for HA and Bursicon-α. Note that Bursicon-α expressing somata are enriched in HA-Rab2. Scale bar, 10 µm.

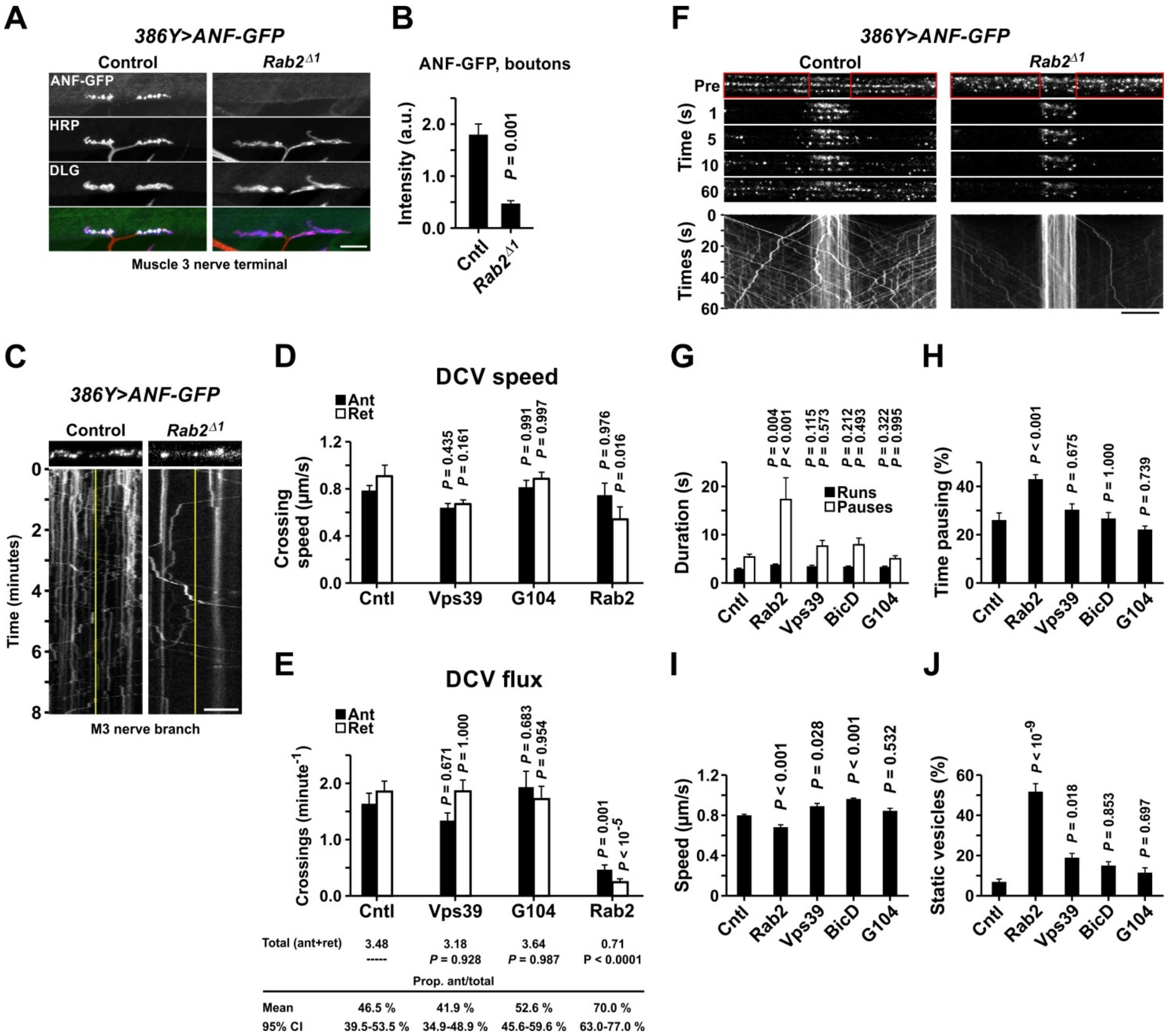


**Figure S5. Impeded axonal transport of ANF-GFP-positive DCVs in *Rab2$^{\Delta1}$* larvae. (A-E)** *Rab2* null mutants exhibit reduced levels of ANF-GFP in muscle 3 type terminals and reduced axonal transport of ANF-GFP-positive DCVs in the associated distal axon. **(A)** *386Y*>ANF-GFP signal and DLG and HRP immunoreactivities in 1b boutons contacting muscle 3. **(B)** Intensity of *386Y*>ANF-GFP in the boutons in **(A)**. Data from four control and four *Rab2$^{\Delta1}$* larvae. Student's *t*-test. **(C)** Kymographs depicting the transport of ANF-GFP-positive DCVs near the distal branch point of the muscle 3 nerve. The first image in the time-lapse series underlying the kymograph is shown at the *top*. The speed and flux (mean ± s.e.m.) of DCVs crossing the *yellow line* in **(C)** is shown in **(D)** and **(E)**, respectively. Ant, anterograde; Ret, retrograde. Each genotype (control, *Vps39$^{\Delta1}$*, *Golgin104$^{ins1}$*, and *Rab2$^{\Delta1}$*) represented by eight larvae. The total flux (anterograde plus retrograde) and the proportion of anterograde crossings (mean 95% ± confidence interval) is shown in **(E)** (*bottom*). *P*-values in **(D and E)** from ANOVAs followed by Tukey's test. **(F-J)** Inhibitory effect of Rab2 loss on axonal transport of ANF-GFP-positive DCVs in mid-axons. **(F)** Time-lapse images and kymographs from control and *Rab2* mutant larvae expressing ANF-GFP widely in peptidergic neurons. Axonal transport of ANF-GFP-positive DCVs in the A8/9 nerve is shown before (Pre), and at indicated time points after bilateral photobleaching (*red boxes*). **(G-J)** Duration of runs and pauses **(G)**, fraction of the duty cycle that the DCVs were pausing **(H)**, run speed **(I)**, and proportion of static vesicles **(J)** in control, *Rab2*$^{\Delta 1}$, *Vps39*$^{\Delta 1}$*/Df*, *BicD*$^{r5}$*/Df*, and *Golgin104*$^{ins1}$ larvae. Data are mean ± s.e.m. Each genotype represented by six preparations. ANOVAs followed by Tukey's test. Scale bars: **(A)**, 20 μm; **(C)**, 5 μm; **(F)**, 10 μm.

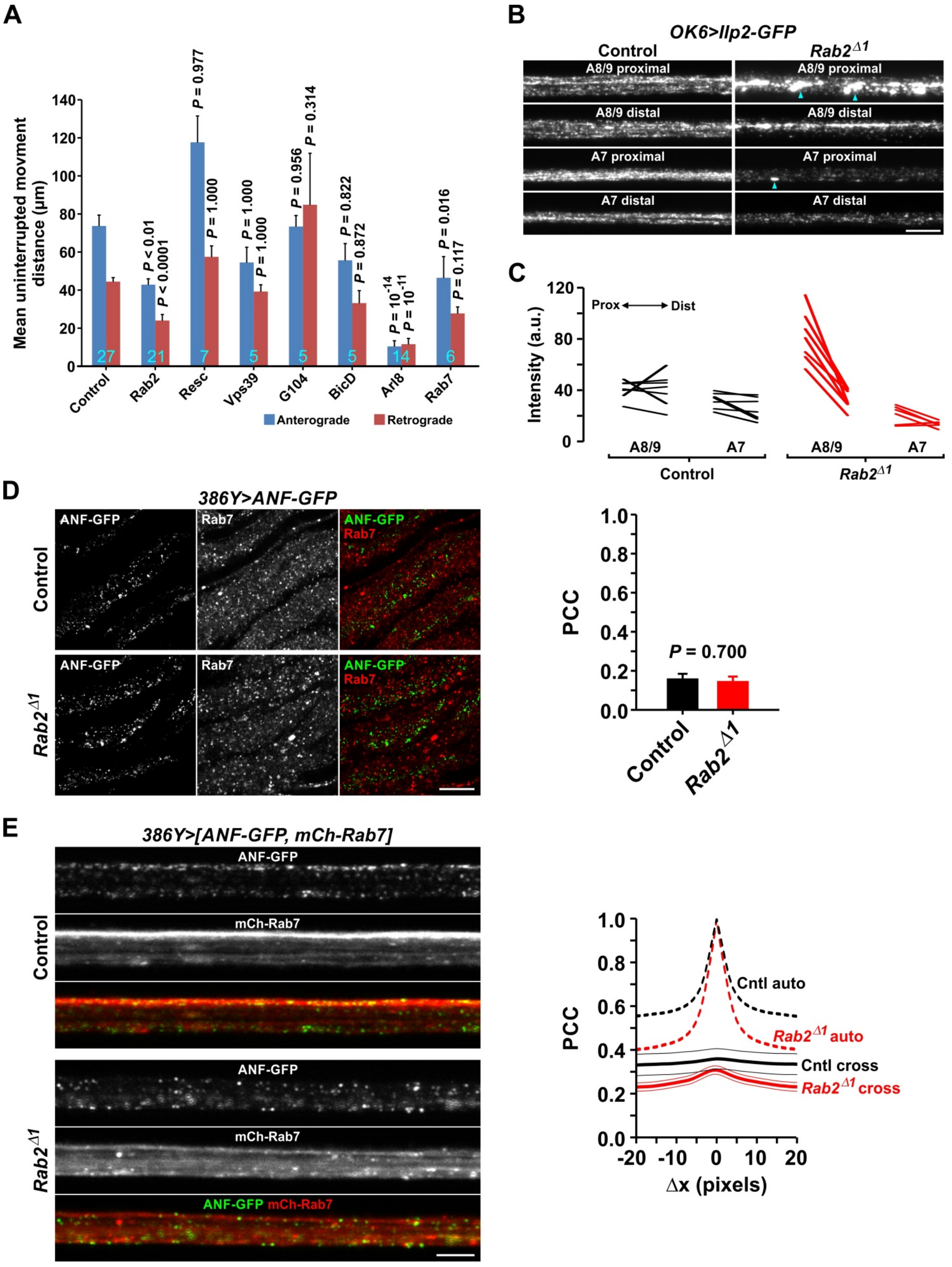
A
Mean uninterrupted movment distance (μm)
Control
Rab2
Resc
Vps39
G104
BicD
Arl8
Rab7
P < 0.01
P < 0.0001
P = 0.977
P = 1.000
P = 1.000
P = 1.000
P = 0.956
P = 0.314
P = 0.822
P = 0.872
P = 10-14
P = 10-11
P = 0.016
P = 0.117
Anterograde
Retrograde
B
OK6>Ilp2-GFP
Control
Rab2Δ1
A8/9 proximal
A8/9 distal
A7 proximal
A7 distal
C
Intensity (a.u.)
Prox
Dist
A8/9
A7
Control
Rab2Δ1
D
386Y>ANF-GFP
ANF-GFP
Rab7
Control
Rab2Δ1
PCC
P = 0.700
E
386Y>[ANF-GFP, mCh-Rab7]
ANF-GFP
mCh-Rab7
Cntl auto
Rab2Δ1 auto
Cntl cross
Rab2Δ1 cross
Δx (pixels)

**Figure S6. Processivity of Ilp2-GFP-positive DCVs, distribution of Ilp2-GFP signal in motor axons along the proximo-distal axis, and absence of substantial overlap between ANF-GFP-positive DCVs and the late endosomal marker Rab7. (A)** Estimated mean uninterrupted movement distance for Ilp2-GFP-positive DCVs, derived from the data in (Fig. 4) by dividing the total observed run distance by the total number of pauses. Data are mean ± s.e.m. The number of larvae analyzed is indicated *inside bars* for each genotype. ANOVA followed by Dunnett's test. **(B)** Sum projections of proximal and distal A8/9 and A7 nerve segments in larvae expressing Ilp2-GFP in motor neurons. *Arrowheads* indicate Ilp2-GFP-dense aggregates. **(C)** Quantification of the signal intensities in **(B)**. Data from seven control and seven *Rab2*$^{\Delta 1}$ larvae. A.u., arbitrary units. **(D)** (*left*) Relationship between ANF-GFP and the endogenous Rab7 immunosignal in the peripheral nerves of control and *Rab2*$^{\Delta 1}$ larvae. **(D)** (*right*) Quantification of **(D)** (*left*). Pearson's cross correlation (PCC) between the ANF-GFP and Rab7 signals is shown. Data (mean ± s.e.m.) from four control and six *Rab2*$^{\Delta 1}$ preparations. **(E)** (*left*) Relationship between ANF-GFP and mCherry-tagged Rab7 in the A8/9 nerve (live imaging). See also Video 2. **(E)** (*right*) Quantification of **(E)** (*left*). Pearson's cross-correlation (cross), indicated with *thick* (mean) and *thin* (± s.e.m.) *solid lines*, is shown as a function of the movement of the GFP and mCherry channels relative to each other (Δx; van Steensel analysis). For each genotype, the autocorrelations (auto) of both the GFP and the mCherry signals were also calculated as a function of Δx, and their average included for reference (*dashed lines*). Data from three control and six *Rab2*$^{\Delta 1}$ preparations. Scale bars: **(B)**, 10 µm; **(D)**, 10 µm; **(E)**, 5 µm.

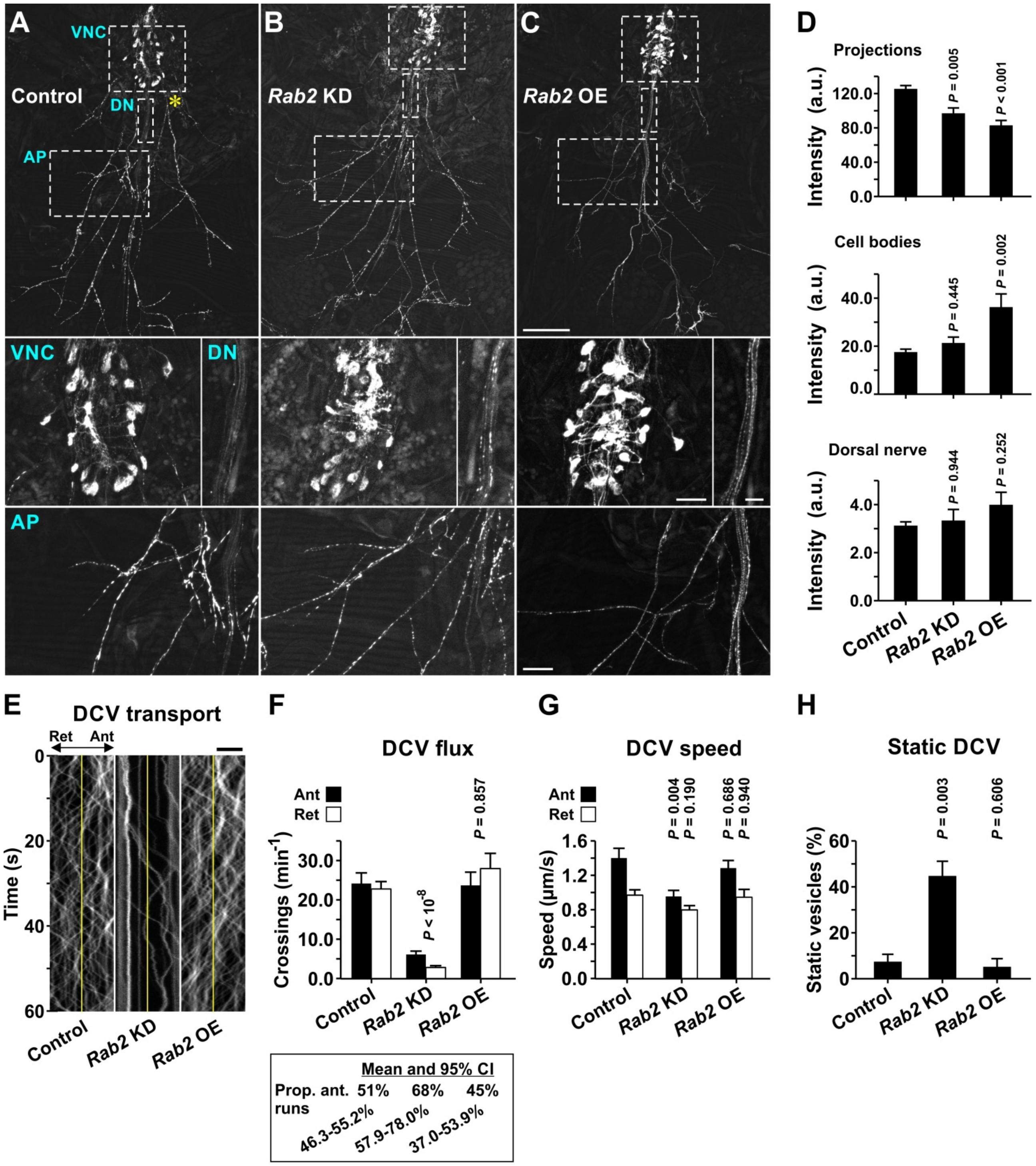


**Figure S7. Effects of *Rab2* depletion and overexpression in pharate adults. (A-C)** Confocal sum projections of *CCAP*-Gal4-directed Ilp2-GFP expression in Bursicon-secreting neurons in fixed pharate adult preparations. **(A)**, Control; **(B)**, *Rab2* knockdown; **(C)**, *Rab2* overexpression driven by *CCAP*-Gal4. Details of the Ilp2-GFP signal in cell bodies in the ventral nerve cord (VNC), the dorsal nerve (DN), and the axonal projections rich in boutons en passant (AP) are shown. **(D)** Quantification of Ilp2 signal intensities in **(A-C)**. Data from six controls, four *Rab2*-depleted, and four *Rab2*-overexpressing animals. **(E-H)** Analysis of the transport of Ilp2-GFP-positive DCVs in the axonal shaft at a position corresponding to the *asterisk* in **(A)**. Data from nine controls, six *Rab2*-depleted, and seven *Rab2*-overexpressing animals. **(E)** Kymographs derived from time-lapse imaging of a 10 µm wide axonal segment. Ant, anterograde; Ret, retrograde. Number **(F)** (*top*) and speed **(G)** of DCVs crossing the *yellow line* in **(E)** during one minute. **(F)** (*bottom*) Proportion (mean ± 95% confidence interval) of anterograde crossings relative to the total number of crossings. The *P*-values at **(F)** (*top*) refers to the total number of crossings, irrespective of direction. **(H)** Proportion of static vesicles within the field of view in **(E)**. ANOVA followed by Tukey's test in **(D)** and **(F-H)**. Scale bars: **(C)** *top* 200 µm; *middle left*, 50 µm; *middle right*, 20 µm; **(C)** *bottom*, 50 µm; **(E),** 4 µm. All data in bar graphs are mean ± s.e.m.

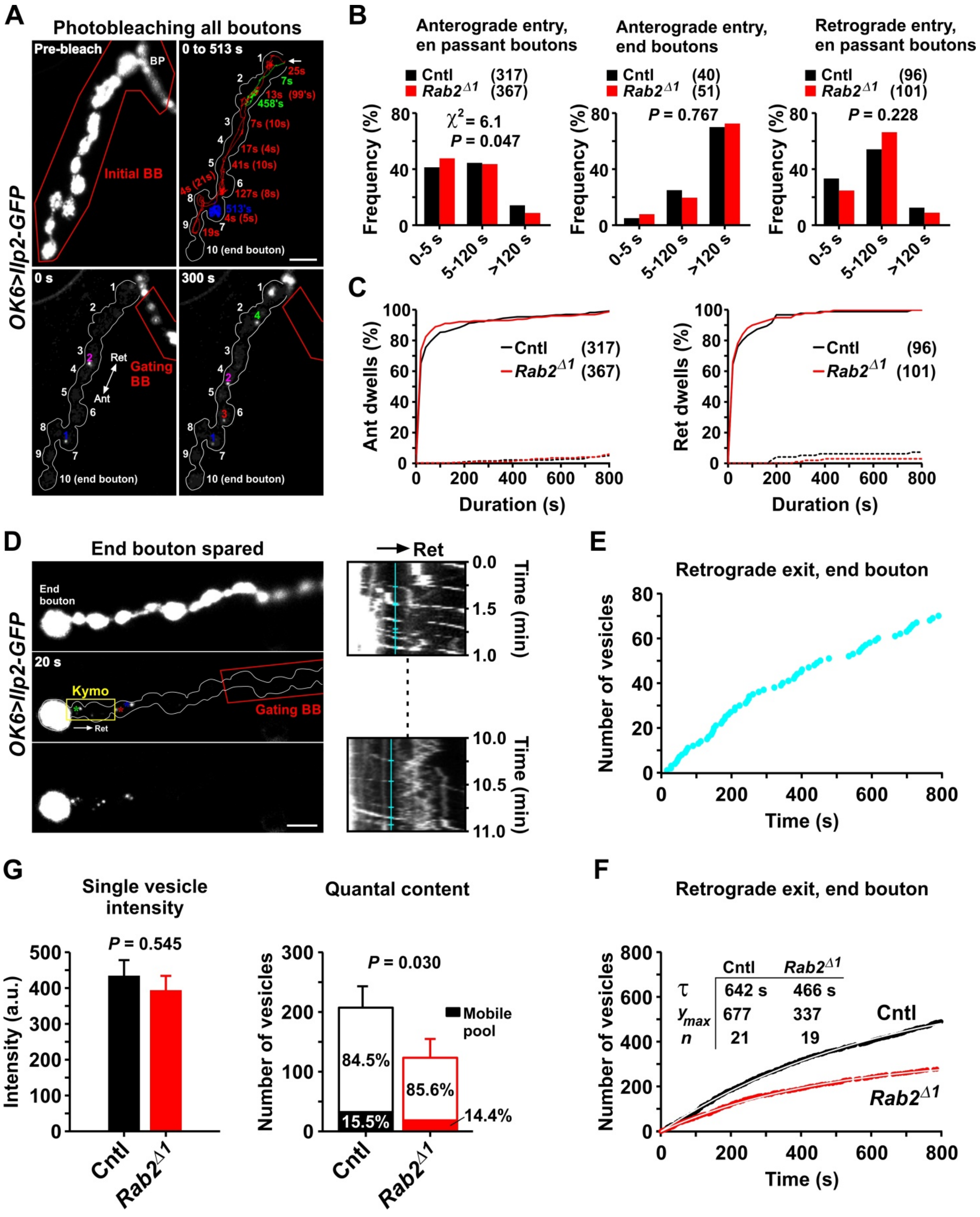


**Figure S8. Transport of Ilp2-GFP-labeled DCVs in larval motor nerve terminals innervating muscle 3. (A)** Time-lapse imaging during a 513 s period following photobleaching of all boutons including the branch point (BP) region (bleach box at *top left*) in control larva. **(A)** (*bottom*) Two vesicles (labeled 1, 2) enter the terminal soon after bleaching and are joined by two more vesicles (3, 4) during the next 300 s. Further entry of fluorescent vesicles was blocked by intermittently bleaching the indicated gating bleach box. **(A)** (*top right*) Trajectories of vesicle 1, 3, and 4 (2 not included for clarity). Vesicle 1 remains in bouton 7; vesicle 3, after entering bouton 1 (*arrow*), travels anterogradely to the second-last bouton 9, then retrogradely to bouton 2; vesicle 4 proceeds to bouton 2 upon entering bouton 1. *Colored numbers* indicate how long the vesicle resides in a bouton (dwell time) after reaching it either in the anterograde (*numbers without parentheses*) or retrograde (*numbers in parentheses*) direction. Censored dwell times are marked by an *apostrophe*. **(B and C)** Frequency distributions of dwell times in control and *Rab2*$^{\Delta 1}$ mutants. *Dashed lines* in **(C)** represent censored values. Numbers of analyzed events are shown in parentheses. **(D)** (*left*) Imaging in control larva after photobleaching all except the end bouton, showing the post-bleach retrograde exit of three vesicles from the end bouton at 20 s (*blue, red, and green asterisks*), supplemented by a few more at 30 s (*yellow asterisks*). Anterograde traffic was eliminated by intermittently bleaching the indicated gating bleach box covering the proximal terminal. **(D)** (*right*) Excerpts (0 and 10 min) of kymograph derived from the region just proximal to the end bouton (*yellow boxes*). **(E)** End bouton exit times extracted from the kymograph in **(D)**. **(F)** Exit times pooled from all end boutons (*n* indicated in *table inset*) in control and *Rab2*$^{\Delta 1}$ mutants. **(G)** (*left*) Mean intensity of exited single vesicles. **(G)** (*right*) Pre-bleach quantal content of end boutons, estimated by dividing their total intensity by the mean single vesicle intensity. Proportions of immobile and mobile vesicles, estimated from the exponential decay fit in **(F)**, are indicated. Number of boutons (larvae) analyzed in **(F)** and **(G)**: Cntl, 21(18); *Rab2*$^{\Delta 1}$, 19(15). Data in **(F)** and **(G)** are mean ± s.e.m. A.u., arbitrary units. Scale bars: 5 µm.

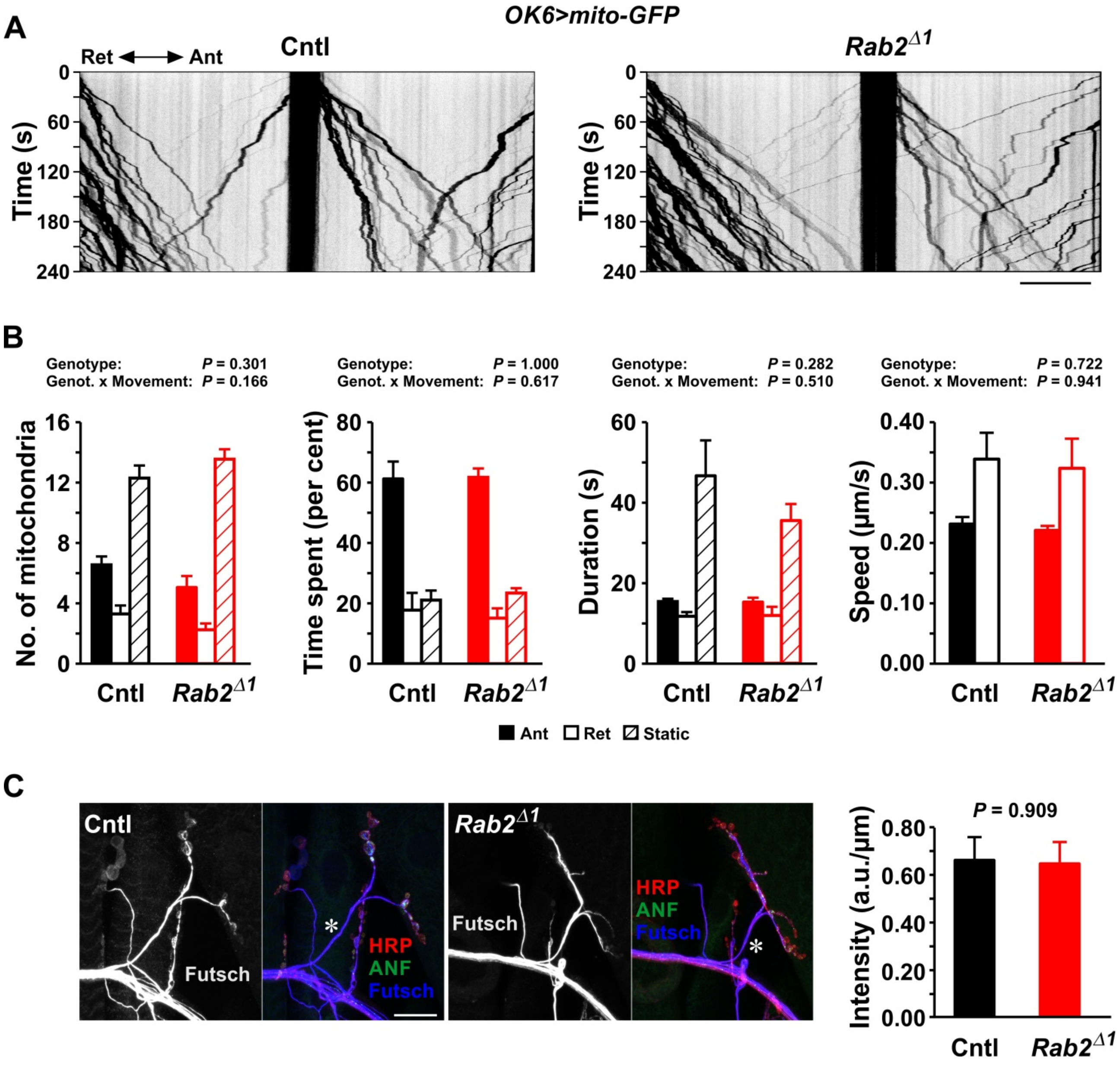


**Figure S9. Loss of Rab2 does neither affect axonal transport of mitochondria, nor the distribution of the neuronal microtubule marker Futsch. (A)** Kymographs (inverted images) showing transport of mito-GFP-labeled mitochondria in motor axons of the A7 peripheral nerve. Lateral regions photobleached at the start of the recording as in (Fig. 3) to enhance the visibility of organelles entering from unbleached areas. **(B)** Kinetic analysis of mitochondrial traffic. From left to right (mean ± s.e.m.): counts of static and dynamic mitochondria, duty cycle, mean duration of individual kinetic events, and run speed. Two-factor MANOVAs. Data from ten control and ten *Rab2*$^{\Delta 1}$ mutant larvae. **(C)** (*left*) Sum projections showing the axon (*asterisk*) projecting to the muscle 3 NMJ in control and *Rab2* null larvae. The Futsch immunosignal is shown separately or merged with HRP immunosignal and raw ANF-GFP fluorescence. **(C)** (*right*) Futsch intensity (mean ± s.e.m.) in the distal axon innervating muscle 3, measured in the approximate region marked by the asterisk in **(C)** (left) and corrected for axon length. Student's *t*-test ($n$ = four preparations for both genotypes). A.u., arbitrary units. Scale bars: **(A)**, 10 µm; **(B)**, 20 µm.

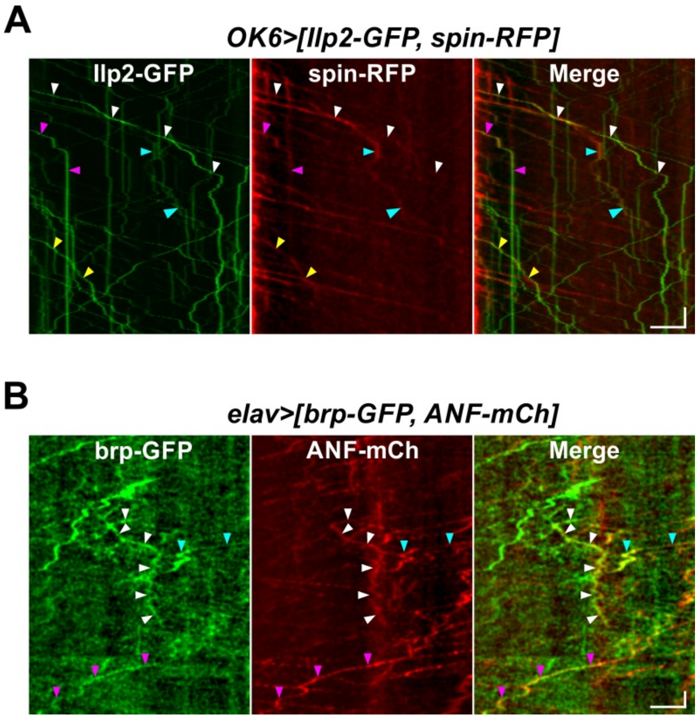


**Figure S10. Examples of co-transport of DCV cargo and lysosomal-related vesicle markers.** Dual-channel kymographs showing co-transport events of DCV markers (Ilp2-GFP and ANF-mCherry) with the lysosomal marker spin-RFP **(A)** and the active zone marker brp-GFP **(B)** in peripheral nerve axons. *Arrowheads* indicate overlapping tracks. Note that most tracks do not overlap. The red channel in **(A)** and both channels in **(B)** were band-pass filtered to enhance visibility (see Methods). Segmental nerves in L3 larvae, live confocal time series. Nerves were photobleached prior to recording of the underlying time series to eliminate fluorescence from static structures. Images were selected from the following number of experiments (larvae): **(A)**, 6 (1): **(B)**, 28 (8). Scale bars (horizontal): **(A)**, 10 s; **(B)**, 20 s. Scale bars (vertical), 5 µm.

**Table S1.** Detailed genotypes. *w*$^{1118}$ served as wild type.

| Figure or Video | Panel and genotypes |
|---|---|
| **Fig. 1** | **A, B, C**<br>***HA-Rab2/c929-Gal4 UAS-GFP-Rab11***<br><br>**D**<br>***HA-Rab2***<br><br>**E, F**<br>+<br>***Rab2***$^{\Delta1}$<br>***Rab2***$^{\Delta1}$***; Tub-Gal4 UAS-mCherry-Rab2/+***<br><br>**G**<br>+<br>***Rab2***$^{\Delta1}$ |
| **Fig. 2** | **A, B**<br>+<br>***Rab2***$^{\Delta1}$<br><br>**C-H**<br>***386Y-Gal4 UAS-ANF-GFP/+***<br>***Rab2***$^{\Delta1}$***; 386Y-Gal4 UAS-ANF-GFP/+*** |
| **Fig. 3** | **A**<br>***OK6-Gal4 UAS-Ilp2-GFP/+***<br>***Rab2***$^{\Delta1}$ ***OK6-Gal4/Rab2***$^{\Delta1}$ ***UAS-Ilp2-GFP***<br><br>**B-F**<br>***OK6-Gal4 UAS-Ilp2-GFP/+***<br>***Rab2***$^{\Delta1}$ ***OK6-Gal4/Rab2***$^{\Delta1}$ ***UAS-Ilp2-GFP***<br>***Rab2***$^{\Delta1}$ ***OK6-Gal4/Rab2***$^{\Delta1}$ ***UAS-Ilp2-GFP; UAS-Rab2*** $^{RNAi\text{-}insens(1)}$***/+***<br>***OK6-Gal4 UAS-Ilp2-GFP/+; Vps39*** $^{\Delta1}$***/Df(3R)Exel6178 P{w[+mC]=XP-U}Exel6178***<br>***OK6-Gal4 UAS-Ilp2-GFP/+; Golgin104*** $^{ins1}$***/Df(3L)BSC649***<br>***BicD*** $^{\Delta1}$ ***OK6-Gal4 UAS-Ilp2-GFP/Df(2L)Exel7068***<br>***OK6-Gal4 UAS-Ilp2-GFP/+; Arl8*** $^{e00336}$***/Df(3R)BSC222***<br>***OK6-Gal4 UAS-Ilp2-GFP/+; Rab7*** $^{\Delta1}$ |
| **Fig. 4** | **A-E**<br>***OK6-Gal4 UAS-Ilp2-GFP/+***<br>***Rab2***$^{\Delta1}$ ***OK6-Gal4/ Rab2***$^{\Delta1}$ ***UAS-Ilp2-GFP*** |
| **Fig. 5** | **A**<br>***OK6-Gal4/+; UAS-Rab5-GFP/+***<br>***Rab2***$^{\Delta1}$ ***OK6-Gal4/Rab2***$^{\Delta1}$***; UAS-Rab5-GFP/+***<br><br>**B**<br>***OK6-Gal4/+; UAS-Rab7-GFP/+***<br>***Rab2***$^{\Delta1}$ ***OK6-Gal4/Rab2***$^{\Delta1}$***; UAS-Rab7-GFP/+*** |

| | |
|---|---|
| | **C**<br>*OK6-Gal4/+; UAS-spin.myc-GFP/+*<br>*Rab2$^{\Delta 1}$ OK6-Gal4/Rab2$^{\Delta 1}$; UAS-spin.myc-GFP/+*<br><br>**D**<br>*OK6-Gal4/+; UAS-brp-GFP/+*<br>*Rab2$^{\Delta 1}$ OK6-Gal4/Rab2$^{\Delta 1}$; UAS-brp-GFP/+* |
| **Fig. 6** | **A**<br>*GFP-Rab2*<br>*OK6-Gal4/UAS-pHlourin-Rab2$^{S20N}$*<br>*OK6-Gal4/UAS-pHlourin-Rab2$^{Q65L}$*<br><br>**B**<br>*elav-Gal4/+(/Y); GFP-Rab2/+; UAS-ANF-mCherry/+*<br>*elav-Gal4/+(/Y); UAS-pHlourin-Rab2/+; UAS-ANF-mCherry/+*<br>*elav-Gal4/+(/Y); UAS-pHlourin-Rab2$^{Q65L}$/+; UAS-ANF-mCherry/+*<br><br>**C**<br>*elav-Gal4/+(/Y);; UAS-2xHA-TurboID-Rab2/+*<br>*elav-Gal4/+(/Y);; UAS-2xHA-TurboID-Rab2$^{S20N}$/+*<br>*elav-Gal4/+(/Y);; UAS-2xHA-TurboID-Rab2$^{Q65L}$/+*<br>+ |
| **Fig. 7** | **A**<br>*OK6-Gal4 UAS-Ilp2-GFP/+*<br>*OK6-Gal4 UAS-Ilp2-GFP/+; Arl8$^{e00336}$/Df(3R)BSC222*<br><br>**B**<br>*OK6-Gal4 UAS-Ilp2-GFP/+*<br>*OK6-Gal4 UAS-Ilp2-GFP/+; Arl8$^{e00336}$/Df(3R)BSC222*<br>*Rab2$^{\Delta 1}$ OK6-Gal4/Rab2$^{\Delta 1}$ UAS-Ilp2-GFP*<br><br>**C**<br>*elav-Gal4/+(/Y); UAS-Arl8-GFP/+; UAS-ANF-mCherry/+* |
| **Fig. S1** | **A-C**<br>*UAS-DCR2/+;; 386Y-Gal4 UAS-mCD8-GFP/+*<br>*UAS-DCR2/+; UAS-Rab2-RNAi$^{KK107630}$/+; 386Y-Gal4 UAS-mCD8-GFP/+* |
| **Fig. S2** | **A-C**<br>*386Y-Gal4 UAS-ANF-GFP/+*<br>*Rab2$^{\Delta 1}$; 386Y-Gal4 UAS-ANF-GFP/+*<br><br>**D-F**<br>*386Y-Gal4 UAS-IA2-GFP/+*<br>*Rab2$^{\Delta 1}$; 386Y-Gal4 UAS-IA2-GFP/+* |
| **Fig. S3** | **A, B**<br>*386Y-Gal4 UAS-ANF-GFP/+*<br>*Rab2$^{\Delta 1}$; 386Y-Gal4 UAS-ANF-GFP/+*<br><br>**C**<br>*386Y-Gal4 UAS-ANF-GFP/UAS-Rab2$^{RNAi\text{-}insens(1)}$*<br>*386Y-Gal4 UAS-ANF-GFP/+*<br>*Rab2$^{\Delta 1}$; 386Y-Gal4 UAS-ANF-GFP/+*<br>*Vps39$^{\Delta 1}$ 386Y-Gal4 UAS-ANF-GFP/Vps39$^{\Delta 1}$* |

| | |
|---|---|
| | *Golgin104$^{ins1}$ 386Y-Gal4 UAS-ANF-GFP/Golgin104$^{ins1}$*<br>*CG3703$^{\Delta 1}$ 386Y-Gal4 UAS-ANF-GFP/CG3703$^{\Delta 1}$*<br>*PICK1$^{1}$/PICK1$^{2}$; 386Y-Gal4 UAS-ANF-GFP/+*<br>*BicD$^{r5}$/Df(2L)Exel7068; 386Y-Gal4 UAS-ANF-GFP/+*<br><br>**D**<br>*+*<br>*Rab2$^{\Delta 1}$*<br>*Vps39$^{\Delta 1}$*<br>*Golgin104$^{ins1}$*<br>*Golgin45$^{\Delta 1}$*<br>*CG3703$^{\Delta 1}$*<br>*PICK1$^{\Delta 1}$*<br>*BicD$^{r5}$/Df(2L)Exel7068*<br>*lt$^{11}$/Df(2L)lt45 PPO1[Bc]*<br>*dor$^{8}$*<br><br>**E, F**<br>*386Y-Gal4 UAS-ANF-GFP/+*<br>*Rab2$^{\Delta 1}$; 386Y-Gal4 UAS-ANF-GFP/+*<br><br>**G**<br>*HA-Rab2/+; 386Y-Gal4 UAS-Golgin245-GFP/+* |
| **Fig. S4** | **A**<br>*UAS-DCR2/+; CCAP-Gal4/+*<br>*UAS-DCR2/+; CCAP-Gal4/UAS-Rab2-RNAi$^{KK107630}$*<br><br>**B**<br>*UAS-DCR2/+; UAS-Rab2-RNAi$^{KK107630}$/+*<br>*UAS-DCR2/+; UAS-Rab2-RNAi$^{KK107630}$/OK371-Gal4*<br>*UAS-DCR2/+; UAS-Rab2-RNAi$^{KK107630}$/Gad1-Gal4*<br>*UAS-DCR2/+; UAS-Rab2-RNAi$^{KK107630}$/ChAT-Gal4*<br>*UAS-DCR2/+; UAS-Rab2-RNAi$^{KK107630}$/+; ple-Gal4/+*<br>*UAS-DCR2/+; UAS-Rab2-RNAi$^{KK107630}$/Ddc-Gal4*<br>*UAS-DCR2/+; UAS-Rab2-RNAi$^{KK107630}$/c929-Gal4*<br>*UAS-DCR2/+; UAS-Rab2-RNAi$^{KK107630}$/+; 386Y-Gal4/+*<br>*UAS-DCR2/+; UAS-Rab2-RNAi$^{KK107630}$/CCAP-Gal4*<br>*UAS-DCR2/+; UAS-Rab2-RNAi$^{KK107630}$/+; Ilp2-Gal4/+*<br><br>**C**<br>*UAS-DCR2/+; Burs-Gal4/+*<br>*UAS-DCR2/+; UAS-Rab2-RNAi$^{KK107630}$/Burs-Gal4*<br>*UAS-DCR2/+; CCAP-Gal4 UAS-Rab2-RNAi$^{KK107630}$/+*<br>*UAS-DCR2/+; CCAP-Gal4 UAS-Rab2-RNAi$^{KK107630}$/+; UAS-Rab2$^{RNAi\text{-}insens(1)}$/+*<br>*UAS-DCR2/+; CCAP-Gal4 UAS-Rab2-RNAi$^{KK107630}$/+; UAS-Rab2$^{RNAi\text{-}insens(2)}$/+*<br>*UAS-DCR2/+; CCAP-Gal4 UAS-Rab2-RNAi$^{KK107630}$/UAS-mCherry-Rab2*<br>*UAS-DCR2/+; CCAP-Gal4 UAS-Rab2-RNAi$^{KK107630}$/+; UAS-mCherry-Rab2/+*<br>*UAS-DCR2/+; CCAP-Gal4 UAS-Rab2-RNAi$^{KK107630}$/UAS-Rab11-GFP*<br>*UAS-DCR2/+; CCAP-Gal4 UAS-Rab2-RNAi$^{KK107630}$/+; UAS-IA2-GFP/+* |
| **Fig. S5** | **A-C, F**<br>*386Y-Gal4 UAS-ANF-GFP/+*<br>*Rab2$^{\Delta 1}$; 386Y-Gal4 UAS-ANF-GFP/+* |

| | |
|---|---|
| | **D-E**<br>386Y-Gal4 UAS-ANF-GFP/+<br>Vps39$^{\Delta 1}$ 386Y-Gal4 UAS-ANF-GFP/Vps39$^{\Delta 1}$<br>Golgin104$^{ins1}$ 386Y-Gal4 UAS-ANF-GFP/Golgin104$^{ins1}$<br>Rab2$^{\Delta 1}$; 386Y-Gal4 UAS-ANF-GFP/+<br><br>**G-J**<br>386Y-Gal4 UAS-ANF-GFP/+<br>Rab2$^{\Delta 1}$; 386Y-Gal4 UAS-ANF-GFP/+<br>Vps39$^{\Delta 1}$ 386Y-Gal4 UAS-ANF-GFP/Df(3R)Exel6178 P{w[+mC]=XP-U}Exel6178<br>BicD$^{r5}$/Df(2L)Exel7068; 386Y-Gal4 UAS-ANF-GFP/+<br>Golgin104$^{ins1}$ 386Y-Gal4 UAS-ANF-GFP/ Golgin104$^{ins1}$ |
| **Fig. S6** | **A**<br>OK6-Gal4 UAS-Ilp2-GFP/+<br>Rab2$^{\Delta 1}$ OK6-Gal4/Rab2$^{\Delta 1}$ UAS-Ilp2-GFP<br>Rab2$^{\Delta 1}$ OK6-Gal4/Rab2$^{\Delta 1}$ UAS-Ilp2-GFP; UAS-Rab2$^{RNAi\text{-}insens(1)}$/+.<br>OK6-Gal4 UAS-Ilp2-GFP/+; Vps39$^{\Delta 1}$/Df(3R)Exel6178 P{w[+mC]=XP-U}Exel6178<br>OK6-Gal4 UAS-Ilp2-GFP/+; Golgin104$^{ins1}$/Df(3L)BSC649<br>BicD$^{\Delta 1}$ OK6-Gal4 UAS-Ilp2-GFP/Df(2L)Exel7068<br>OK6-Gal4 UAS-Ilp2-GFP/+; Arl8$^{e00336}$/Df(3R)BSC222<br>OK6-Gal4 UAS-Ilp2-GFP/+; Rab7$^{\Delta 1}$<br><br>**B, C**<br>OK6-Gal4 UAS-Ilp2-GFP/+<br>Rab2$^{\Delta 1}$ OK6-Gal4/Rab2$^{\Delta 1}$ UAS-Ilp2-GFP<br><br>**D**<br>386Y-Gal4 UAS-ANF-GFP/+<br>Rab2$^{\Delta 1}$; 386Y-Gal4 UAS-ANF-GFP/+<br><br>**E**<br>386Y-Gal4 UAS-ANF-GFP/UAS-mCherry-Rab7<br>Rab2$^{\Delta 1}$; 386Y-Gal4 UAS-ANF-GFP/UAS-mCherry-Rab7 |
| **Fig. S7** | **A-H**<br>UAS-DCR2/+; CCAP-Gal4 UAS-Ilp2-GFP/+<br>UAS-DCR2/+; CCAP-Gal4 UAS-Ilp2-GFP/UAS-Rab2-RNAi$^{KK107630}$<br>UAS-DCR2/+; CCAP-Gal4 UAS-Ilp2-GFP/+; UAS-Rab2$^{RNAi\text{-}insens(1)}$/+ |
| **Fig. S8** | **A, D, E**<br>OK6-Gal4 UAS-Ilp2-GFP/+<br><br>**B, C, F, G**<br>OK6-Gal4 UAS-Ilp2-GFP/+<br>Rab2$^{\Delta 1}$ OK6-Gal4/Rab2$^{\Delta 1}$ UAS-Ilp2-GFP |
| **Fig. S9** | **A, B**<br>OK6-Gal4 UAS-mito-HA-GFP/+<br>Rab2$^{\Delta 1}$ OK6-Gal4/ Rab2$^{\Delta 1}$ UAS-mito-HA-GFP<br><br>**C**<br>386Y-Gal4 UAS-ANF-GFP/+<br>Rab2$^{\Delta 1}$; 386Y-Gal4 UAS-ANF-GFP/+ |

| | |
|---|---|
| **Fig. S10** | **A**<br>***OK6-Gal4 UAS-Ilp2-GFP/UAS-spin.myc-RFP***<br><br>**B**<br>***elav-Gal4/+(/Y);; UAS-ANF-mCherry/UAS-brp-GFP*** |
| **Video 1** | ***OK6-Gal4 UAS-Ilp2-GFP/+***<br>***$Rab2^{\Delta1}$ OK6-Gal4/$Rab2^{\Delta1}$ UAS-Ilp2-GFP***<br>***$Rab2^{\Delta1}$ OK6-Gal4/$Rab2^{\Delta1}$ UAS-Ilp2-GFP; UAS-Rab2 $^{RNAi\text{-}insens(1)}$/+.*** |
| **Video 2** | ***386Y-Gal4 UAS-ANF-GFP/UAS-mCherry-Rab7***<br>***$Rab2^{\Delta1}$; 386Y-Gal4 UAS-ANF-GFP/UAS-mCherry-Rab7*** |
| **Video 3** | ***OK6-Gal4 UAS-Ilp2-GFP/+***<br>***OK6-Gal4 UAS-Ilp2-GFP/+; Arl8 $^{e00336}$/Df(3R)BSC222***<br>***OK6-Gal4 UAS-Ilp2-GFP/+; Rab7 $^{\Delta1}$*** |
| **Video 4** | ***OK6-Gal4/+; UAS-Rab5-GFP/+***<br>***$Rab2^{\Delta1}$ OK6-Gal4/ $Rab2^{\Delta1}$; UAS-Rab5-GFP/+*** |
| **Video 5** | ***OK6-Gal4/+; UAS-Rab7-GFP/+***<br>***$Rab2^{\Delta1}$ OK6-Gal4/ $Rab2^{\Delta1}$; UAS-Rab7-GFP/+*** |
| **Video 6** | ***OK6-Gal4/+; UAS-spin.myc-GFP/+***<br>***$Rab2^{\Delta1}$ OK6-Gal4/ $Rab2^{\Delta1}$; UAS-spin.myc-GFP/+*** |
